# Abscission Checkpoint Bodies Reveal a New Facet of Abscission Checkpoint Control

**DOI:** 10.1101/2020.10.07.330290

**Authors:** Lauren K. Williams, Douglas R. Mackay, Madeline A. Whitney, Wesley I. Sundquist, Katharine S. Ullman

**Affiliations:** Department of Oncological Sciences, Huntsman Cancer Institute, University of Utah, Salt Lake City, UT 84112, USA; Department of Biochemistry, University of Utah School of Medicine, Salt Lake City, UT 84112-5650, USA

## Abstract

The abscission checkpoint regulates the ESCRT membrane fission machinery and thereby delays cytokinetic abscission to protect genomic integrity in response to residual mitotic errors. The checkpoint is maintained by Aurora B kinase, which phosphorylates multiple targets, including CHMP4C, a regulatory ESCRT-III subunit necessary for this checkpoint. We now report the discovery that cytoplasmic abscission checkpoint bodies (ACBs) containing phospho-Aurora B and tri-phospho-CHMP4C develop in telophase under an active checkpoint. ACBs are derived from Mitotic Interchromatin Granules (MIGs), transient mitotic structures whose components are housed in splicing-related nuclear speckles during interphase. ACB formation requires CHMP4C, and the ESCRT factor ALIX also contributes. ACB formation is conserved across cell types and under multiple circumstances that activate the checkpoint. Finally, ACBs retain a population of ALIX, and their presence correlates with delayed recruitment of ALIX to the midbody where it would normally promote abscission. Thus, a cytoplasmic mechanism helps regulate midbody machinery to delay abscission.

## Introduction

Checkpoints function throughout the cell cycle to ensure accurate and timely coordination of cell cycle events (Hartwell and Weinert, 1989). Late in cell division, when cells remain connected by only a thin intercellular bridge containing a microtubule-rich structure termed the midbody, the abscission checkpoint is active and delays the final cut of cytokinesis if residual mitotic errors are detected (Norden et al., 2006; Steigemann et al., 2009). To date, this checkpoint has been found to be responsive to four conditions: chromatin bridges within the midbody, depletion of particular nuclear pore proteins, tension at the intercellular bridge, and previous DNA replication stress (Lafaurie-Janvore et al., 2013; Mackay et al., 2010; Mackay and Ullman, 2015; Petsalaki and Zachos, 2019; Steigemann et al., 2009). Left unchecked, these errors have the potential to disrupt new daughter cell functions. This is most clear in the case of chromatin bridges, which can break under tension forces, ultimately leading to chromothripsis, a mutational signature commonly found in cancers (Maciejowski et al., 2015; Umbreit et al., 2020). Loss of the abscission checkpoint accelerates abscission and induces genomic instability in cultured cells (Sadler et al., 2018). Thus, the abscission checkpoint appears to protect cells from accumulating damage arising from mitotic errors, which can otherwise promote tumorigenesis.

Once the abscission checkpoint is satisfied, cytokinetic abscission is mediated by the Endosomal Sorting Complexes Required for Transport (ESCRT) pathway, at least in transformed, cultured mammalian cells (Carlton and Martin-Serrano, 2007; Gatta and Carlton, 2019; Morita et al., 2007; Vietri et al., 2020). The ESCRT adaptor protein CEP55 provides an ESCRT recruiting platform (Carlton and Martin-Serrano, 2007; Morita et al., 2007) at a protein-rich structure called the Flemming body (Capalbo et al., 2019; Skop et al., 2004) centrally located within the midbody. CEP55 forms rings on either side of the Flemming body and recruits the early-acting ESCRTs TSG101 (a component of the ESCRT-I complex) and ALIX through a shared binding site (Lee et al., 2008). SEPT9 has also recently been implicated as a second, TSG101/ESCRT-I-specific midbody adaptor (Karasmanis et al., 2019). ALIX and TSG101/ESCRT-I, in turn, ultimately recruit late-acting ESCRT-III factors, including CHMP4B and IST1 through two parallel pathways (Christ et al., 2016). ESCRT-III subunits polymerize into membrane-constricting filaments and interact with the AAA-ATPase VPS4 to sever membranes in abscission zones on either side of the Flemming body (Carlton and Martin-Serrano, 2007; Christ et al., 2016; Morita et al., 2007). The abscission checkpoint must, therefore, inhibit ESCRT recruitment, polymerization or constriction to delay abscission.

Aurora B kinase (**AurB**) functions as the master regulator of the abscission checkpoint. AurB phosphorylates many different targets at the midbody, and thereby enforces abscission delay and stabilizes the midbody (Hegemann et al., 2014; Kettenbach et al., 2011; Steigemann et al., 2009). At mitotic onset, AurB is phosphorylated (**pAurB**) and thus activated, both by the collective action of Clk1, 2, and 4 kinases and by autophosphorylation (Petsalaki and Zachos, 2016; Yasui et al., 2004). When the abscission checkpoint is satisfied, AurB is dephosphorylated and abscission proceeds (Bhowmick et al., 2019). Many pAurB targets likely remain to be elucidated, but one significant target is the regulatory ESCRT-III subunit, CHMP4C, a factor required for abscission checkpoint activity (Capalbo et al., 2012; Carlton et al., 2012). pAurB phosphorylates three sites within a region unique to the CHMP4C isoform that is required for the abscission checkpoint (Figure 1A) (Capalbo et al., 2012; Carlton et al., 2012), and the ESCRT-binding kinase ULK3 phosphorylates other site(s) outside this insert region (Caballe et al., 2015). A naturally occurring variant allele of CHMP4C that reduces binding to ALIX abrogates the abscission checkpoint, induces DNA damage accumulation and correlates with increased susceptibility to several different cancers (Pharoah et al., 2013; Sadler et al., 2018). Thus, CHMP4C phosphorylation and its ALIX binding activity are necessary for abscission checkpoint function.

**Figure 1.**
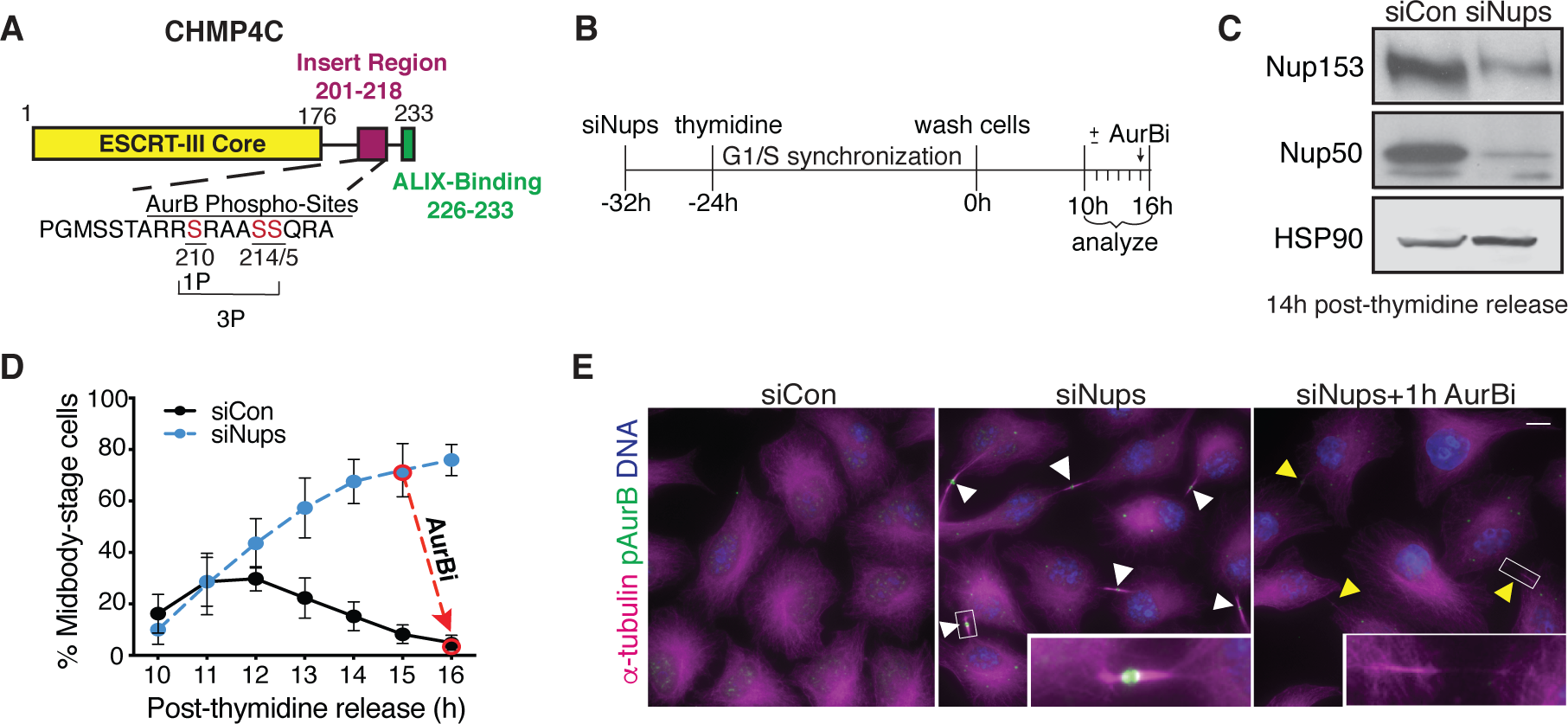
Assay for abscission checkpoint synchronization. (**A**) Schematic of CHMP4C protein domain organization. (**B**) Timeline of abscission checkpoint activation and cell synchronization with siNup153/siNup50 (siNups) or control siRNAs, and thymidine treatment, respectively. (**C**) Western blot of lysates prepared from control (siCon) and checkpoint-activated (siNups) HeLa cells harvested 14 hours post-thymidine release. (**D**) Quantification of checkpoint-activated cells after thymidine release. N=1500 (total) cells/timepoint from n=5 independent biological replicates. (**E)** Images acquired 16 hours post-thymidine release. White arrowheads: midbodies. Yellow arrowheads: abscised midbodies. Scale bar, 10 µm. Insets have enhanced brightness. <u>Throughout manuscript:</u> DNA is detected with DAPI unless noted. Bar and line graphs represent mean ± standard deviation. Times refer to timeline in **B** unless noted. N=total from all replicates.

Checkpoint-induced abscission delay appears to be a multistep process, and previous studies have implicated: 1) phosphoregulation of several ESCRT-III activities (Caballe et al., 2015) and 2) sequestration of VPS4 in a single ring within the Flemming body, away from the abscission zones (Thoresen et al., 2014). In the latter case, phosphorylated CHMP4C acts together with the adaptor protein ANCHR to sequester VPS4 (Thoresen et al., 2014). Yet, the finding that CHMP4C must be able to bind ALIX for the checkpoint to function (Sadler et al., 2018) suggests additional crucial roles for CHMP4C. It remains unknown, however, where such interactions take place and whether cytoplasmic checkpoint regulatory mechanisms complement those at the midbody.

## Results

### Synchronous abscission checkpoint activation

The abscission checkpoint is normally active in only a small fraction of cultured cells because at any given time few cells are in the cell cycle phase of cytokinesis and, further, because checkpoint activity is normally transient. We have overcome these experimental hurdles by generating sustained abscission checkpoint activity in a cell population by combining treatments that: 1) activate the checkpoint (Mackay et al., 2010), using siRNA-mediated depletion of the nuclear pore basket proteins Nup153 and Nup50 (referred to throughout as siNups), and 2) synchronize the cell cycle, using thymidine addition to arrest cells in G1/S phase, followed by release and synchronous progression through the remaining cell cycle (Figure 1B-C). These conditions elicited an active checkpoint in up to ∼80% of HeLa cells in culture (Figure 1D-E). The arrested cells remained competent for abscission, however, as they all completed division within 60 minutes following deactivation of the checkpoint with an AurB inhibitor (**AurBi**, ZM 447439) (Figure 1D-E), or after >5 hours without treatment (not shown).

### Abscission checkpoint activation delays ALIX recruitment to the midbody

Using this new checkpoint induction protocol, we tested whether ESCRT factor recruitment to midbodies was altered by the abscission checkpoint. The ESCRT adaptor CEP55 and one of the two early-acting ESCRT factors, TSG101/ESCRT-I, localized normally to midbodies when the checkpoint was activated (Figure 2A-B, Figure 2—figure supplement 1A, figure supplement 2A-B). In contrast, checkpoint activation induced a significant delay in the midbody recruitment of the other early-acting ESCRT factor, ALIX, with this delay particularly pronounced in earlier-stage midbodies (Figure 2C-D, Figure 2—figure supplement 1B). This result suggested that delayed ALIX recruitment could contribute to the abscission delay dictated by abscission checkpoint activation. To control for potential cell-to-cell variation, we also quantified CEP55 and ALIX intensities within the same cells to determine their relative ratios at individual midbodies. This experiment confirmed that an active checkpoint delays recruitment of ALIX to the midbody relative to CEP55 with an ∼3-fold decrease in the relative ratio 11 hours post-thymidine release (Figure 2E-F, Figure 2—figure supplement 1C), and an ∼3-fold decrease relative to TSG101/ESCRT-I in an analogous experiment (Figure 2—figure supplement 2C-D). By 16 hours after thymidine release, ALIX levels at the midbody had recovered, consistent with the duration of checkpoint signaling under these conditions (Figure 2D, F). AurBi treatment rapidly rescued ALIX recruitment at 11 hours (Figure 2G), confirming that delays in ALIX recruitment were checkpoint dependent. We also found that checkpoint activation delayed recruitment of IST1, an ESCRT-III subunit that functions downstream of ALIX and is required for abscission (Figure 2—figure supplement 3A-B) (Agromayor et al., 2009; Bajorek et al., 2009a). IST1 recruitment was likewise restored by brief AurBi treatment (Figure 2—figure supplement 3C). Taken together, these observations reveal a new dimension of regulation that helps explain why abscission is delayed in response to checkpoint activation. Both ALIX and IST1 localize to midbodies and are required for cytokinesis in HeLa cells (Agromayor et al., 2009; Bajorek et al., 2009a; Carlton and Martin-Serrano, 2007; Morita et al., 2007), and abscission timing is therefore expected to be delayed when their recruitment is delayed.

**Figure 2.**
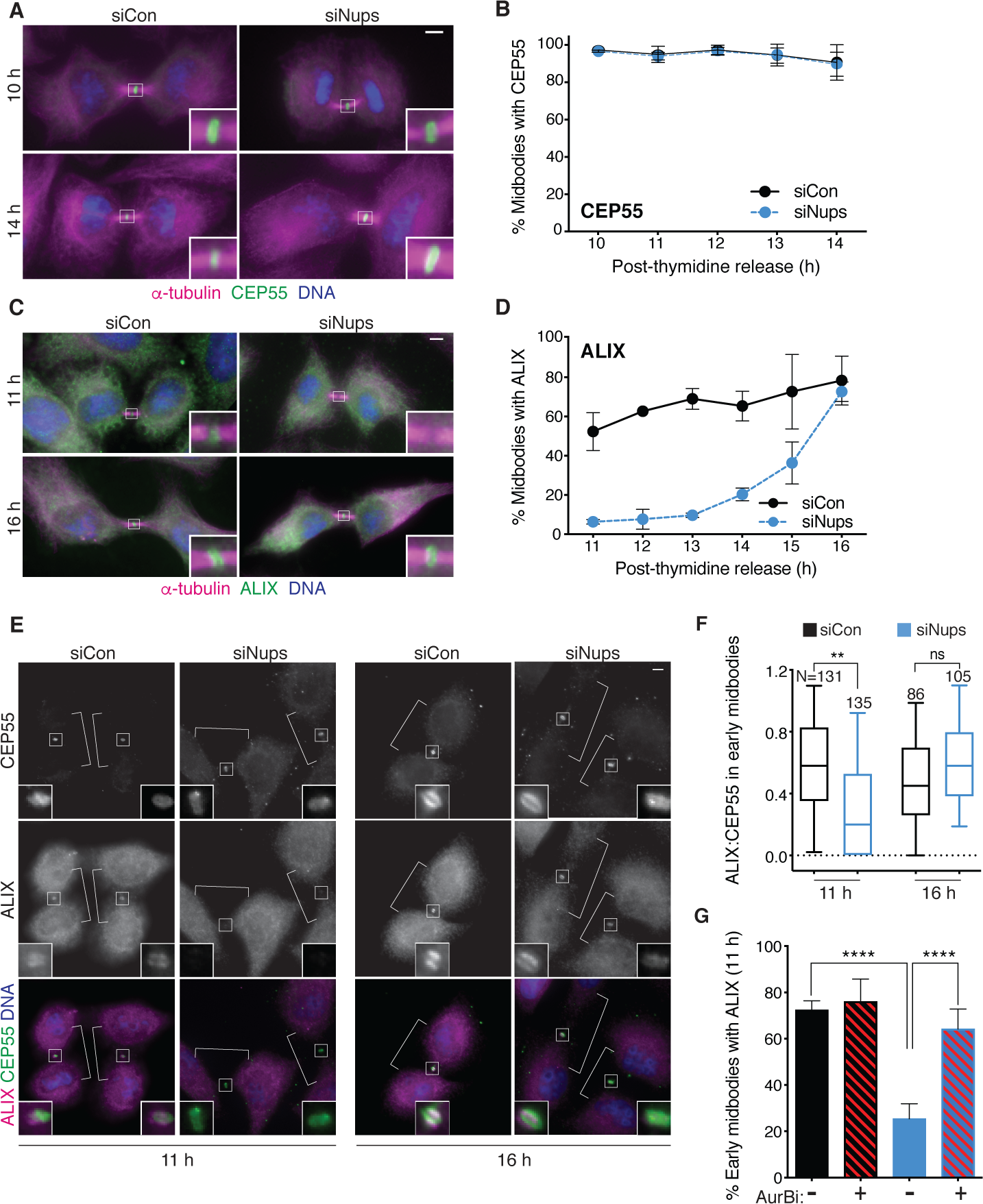
Abscission checkpoint activation delays ALIX recruitment to the midbody. Immunofluorescence and time course quantifications of CEP55 (**A-B**) and ALIX (**C-D**) recruitment to early-stage midbodies in control and checkpoint-activated cells. N=300 midbodies/timepoint from n=3 biological replicates. (**E-F**) Immunofluorescence and quantification of ALIX:CEP55 relative intensities from n=2 biological replicates in individual control and checkpoint-activated early-stage midbodies (see Figure 4B for example of early midbody stage). Only one midbody is shown in the siCon/16 h condition because this sample had fewer midbody-stage cells than other conditions. **(G)** Quantification of ALIX recruitment to early-stage midbodies at 11 h with/without checkpoint activation and with/without AurBi added at 10.5 h. N=300 midbodies/treatment from n=3 biological replicates. <u>Throughout manuscript</u>: scale bars are 5 µm unless noted. White brackets mark midbody-stage cells, as detected by α-tubulin, not shown. Data points without visible error bars (as in 2D siCon 12 h) have a SD too small to display outside the data point or box. Boxplots represent the 25^th^, median, and 75^th^ percentile values. Whiskers represent the 10^th^ and 90^th^ percentiles. P<0.05: *, P≤0.01: **, P≤0.001: ***, P≤0.0001: ****. See methods for statistical tests used.

### Abscission Checkpoint Bodies form in the cytoplasm when the abscission checkpoint is active

We next investigated what other cellular changes occur concomitantly with delayed ALIX recruitment. We have previously shown that CHMP4C mutations that inhibit ALIX binding override the checkpoint (Sadler et al., 2018), indicating that CHMP4C and ALIX bind when the abscission checkpoint is active. We therefore first tested whether CHMP4C localization was also altered when the abscission checkpoint was activated. Stably expressed HA-CHMP4C localized to midbodies regardless of checkpoint status, whereas ALIX recruitment was again delayed when the checkpoint was activated (Figure 3—figure supplement 1A-D). Thus, consistent with previous reports (Sadler et al., 2018), bulk CHMP4C protein localization does not change with checkpoint activation. We next examined the localization of specific AurB-activated phospho-isoforms of CHMP4C using phospho-specific antibodies (Capalbo et al., 2016). As reported previously (Capalbo et al., 2016), singly phosphorylated CHMP4C (pCHMP4C) formed a single ring at the Flemming body and was recruited with the same timing whether or not the checkpoint was activated (Figure 3—figure supplement 1E-F). Tri-phosphorylated CHMP4C (pppCHMP4C) also formed single Flemming body rings and, additionally, formed double rings, one on either side of the Flemming body (Capalbo et al., 2016), particularly when the checkpoint was activated (Figure 3—figure supplement 1G). However, checkpoint-dependent changes in pppCHMP4C distribution were much more dramatic at cytoplasmic sites outside the midbody (Figure 3A-B). In control midbody-stage cells, pppCHMP4C exhibited a granular nuclear localization, whereas checkpoint activation caused pppCHMP4C to accumulate in 0.5-2 µm-diameter cytoplasmic foci that we have named Abscission Checkpoint Bodies (**ACB**s). Pan-CHMP4C antibodies, but not those specific to singly phosphorylated pCHMP4C, decorated ACBs consistent with the tri-phosphorylated subpopulation of CHMP4C preferentially localizing to this site (Figure 3—figure supplement 1E, figure supplement 2A). Note, however, that detection of CHMP4C in ACBs with the pan-antibody required removing soluble cytoplasmic proteins by pre-fixation treatment with buffer (PHEM) to preserve general cell morphology followed by detergent to pre-permeabilize cell membranes. ACBs were observed in the majority of checkpoint-activated cells throughout the entire post-thymidine time course (Figure 3C).

**Figure 3.**
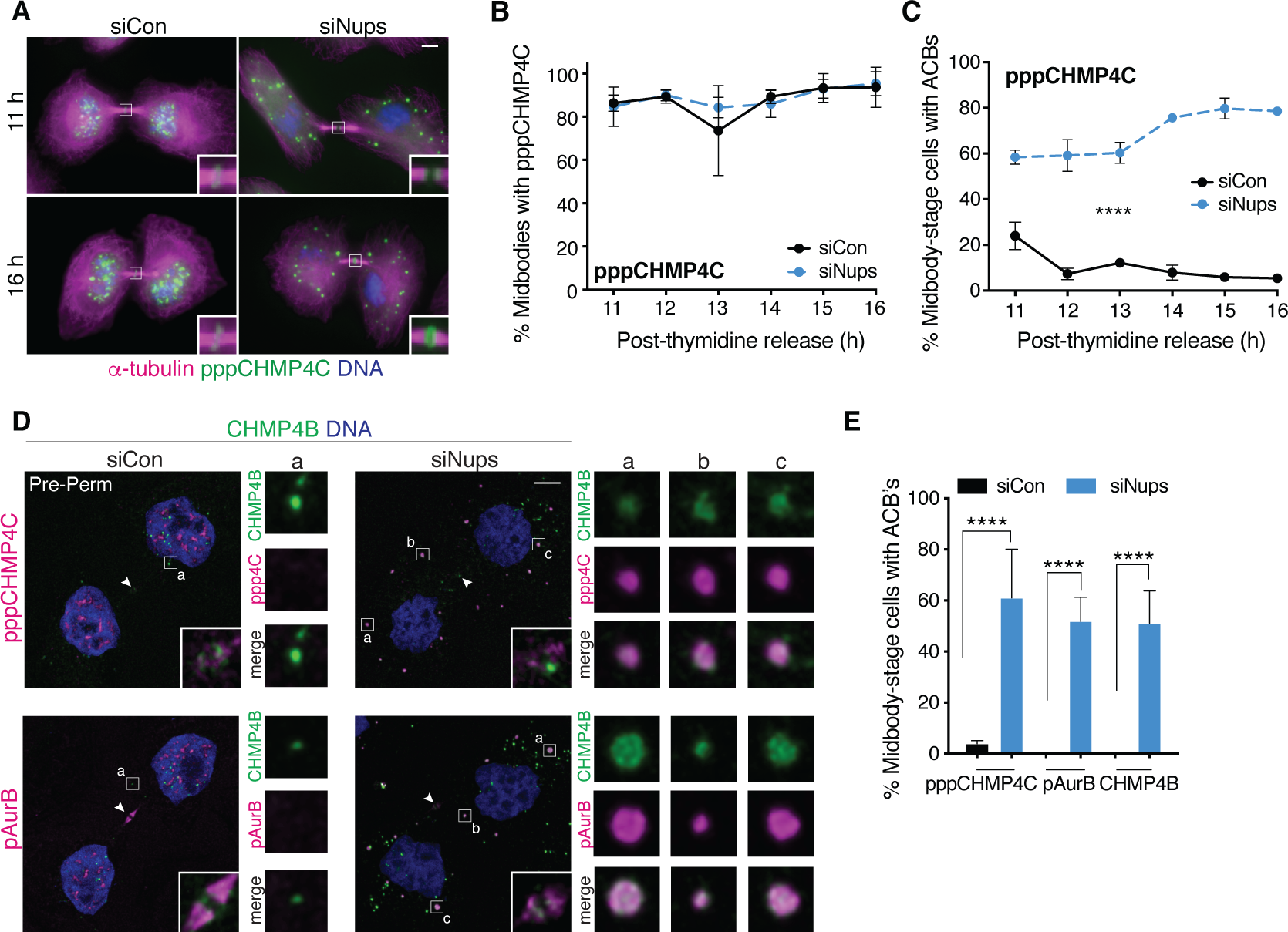
pppCHMP4C localizes to “Abscission Checkpoint Bodies” under an active abscission checkpoint. (**A-B**) Immunofluorescence and time course quantification of pppCHMP4C recruitment to midbodies in control and checkpoint-arrested cells, and (**C**) time course quantification of midbody-stage cells with ACB’s. N ≥ 300 midbodies/timepoint from n=3 biological replicates. (**D**) Confocal z-projections of pre-permeabilized midbody-stage cells under asynchronous conditions (48 h after transfection with siNups or siControl), stained as indicated. Note the appearance of ACB component substructure in some cases. (**E**) Quantification of midbody-stage cells containing ACB’s (treated as in **D**). N=300 midbodies/condition from n=3 biological replicates. <u>Throughout manuscript</u>: Components are brighter in ACBs than in midbodies and the images are optimized for ACBs, which reduces the appearance of midbody localization in the confocal images. Confocal midbody insets are 2.3 µm wide and have enhanced brightness. White arrowheads mark Flemming bodies (where detectable). ACB enlargements are 2 µm wide.

ACBs were abundant (26±4 per midbody-stage cell), and the number of ACBs per midbody-stage cell remained constant throughout the time course (Figure 3—figure supplement 2B). In summary, previously uncharacterized bodies that contain the abscission checkpoint factor pppCHMP4C form in the cytoplasm of cells with an active abscission checkpoint.

### CHMP4B and pAurB also localize to ACBs

Initial screens for additional ACB components revealed that a second CHMP4 isoform, CHMP4B, was also present. These two CHMP4 isoforms have similar sequences, yet perform quite different roles in cell division: CHMP4C regulates abscission timing but is not required for abscission, whereas CHMP4B is required for abscission but does not regulate abscission timing (Capalbo et al., 2012; Carlton et al., 2012, 2008). CHMP4B is an abundant cytosolic ESCRT-III protein and, like bulk CHMP4C, its ACB localization was also best visualized after pre-permeabilizing cells to remove soluble cytoplasmic proteins (Figure 3D). In the absence of checkpoint activation, CHMP4B also exhibited a punctate cytoplasmic distribution in pre-permeabilized cells, but these punctae were significantly smaller than ACBs and did not colocalize with pppCHMP4C. An antibody specific for the T232 phosphorylated, active form of AurB (pAurB) (Yasui et al., 2004) also stained ACBs (Figure 3D), explaining our previous observation that checkpoint activation caused pAurB to localize to cytoplasmic foci of unknown composition (Mackay et al., 2010). Hence, in addition to pppCHMP4C, pAurB and CHMP4B are ACB components and in every case their ACB localization was strongly checkpoint dependent (Figure 3E).

### ACBs are derived from MIGs

We further characterized ACBs by testing whether they corresponded to a variety of known cellular assemblies and organelles (Figure 4—figure supplement S1). Unexpectedly, these experiments revealed that ACBs specifically colocalized with SR domain-containing splicing factors and a phosphorylated form of SC35, both of which are markers of cytoplasmic bodies termed Mitotic Interchromatin Granules (MIGs) (Figure 4A) (Fu and Maniatis, 1990; Li and Bingham, 1991; Reuter et al., 1985). During interphase, MIG components reside in a nuclear compartment called nuclear speckles where these components are hypothesized to be concentrated to increase splicing efficiency (Beck, 1961; Chen and Belmont, 2019). Both MIGs and nuclear speckles are compartments with liquid-liquid phase separation characteristics (Rai et al., 2018; Strom and Brangwynne, 2019) and primarily contain factors that function in mRNA biogenesis, particularly splicing (Mintz et al., 1999; Saitoh et al., 2004; Uversky, 2017). After being released from nuclei upon mitotic nuclear envelope disassembly, these factors associate into small cytoplasmic MIG foci during the metaphase to anaphase transition. MIGs then normally disperse in late telophase, concomitant with step-wise reassembly of nuclear speckles within newly-formed nuclei (Prasanth et al., 2003; Reuter et al., 1985; Saitoh et al., 2012; Spector and Lamond, 2011; Spector and Smith, 1986; Tripathi and Parnaik, 2008). However, abscission checkpoint activation instead promoted the appearance of ACBs, which are larger than MIGs and become apparent in late midbody-stage cells (Figure 4B). Nuclear speckle assembly in the nucleus was delayed concurrently with ACB appearance, as detected by antibodies to pSC35 or the SR domain (Figure 4A-B). We therefore propose that cytokinetic ACBs are derived from MIGs and that abscission checkpoint activation plays a pivotal role in inducing the transition.

**Figure 4.**
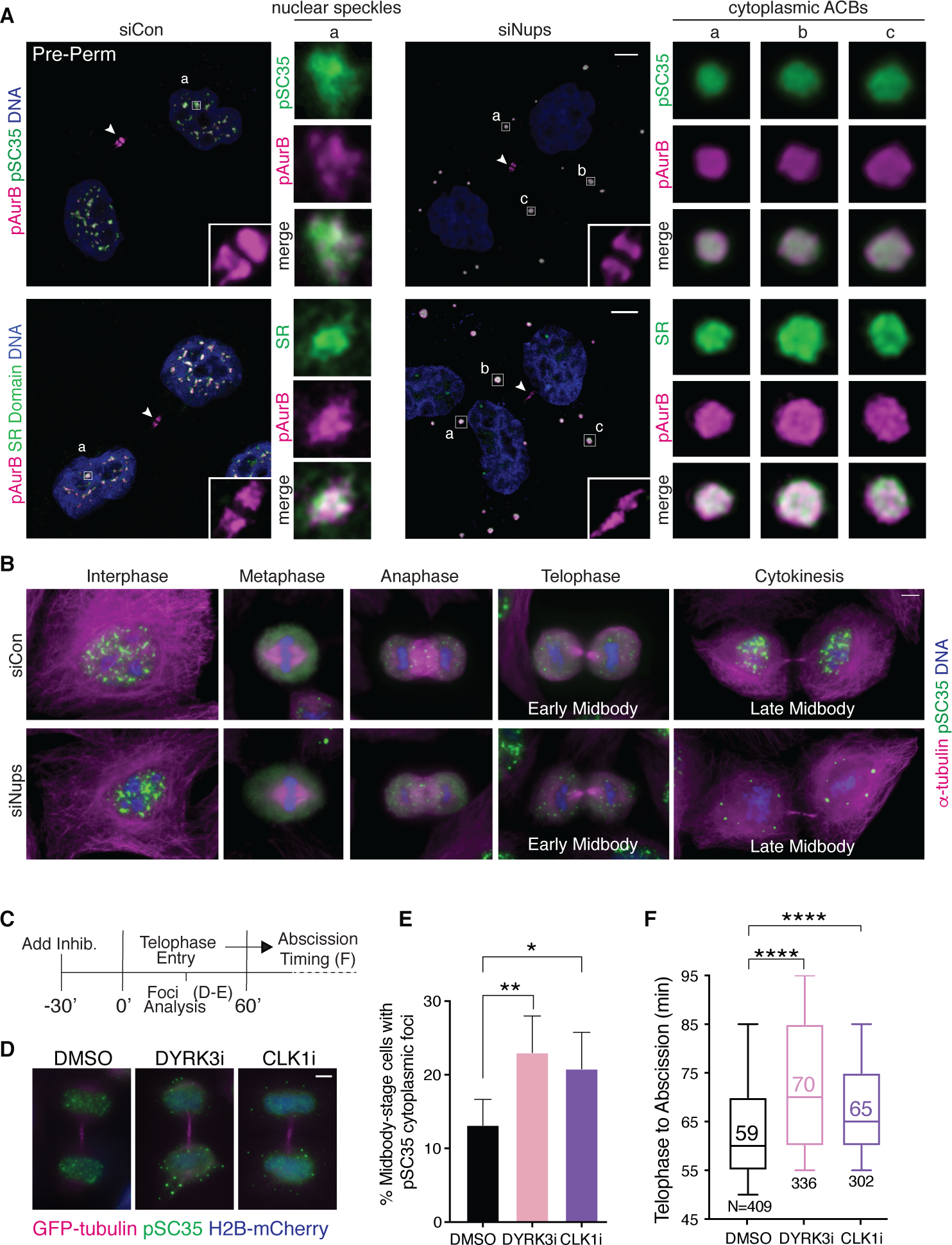
ACBs are related to MIGs and contribute to abscission delay. (**A**) Confocal z-projections of pre-permeabilized (Pre-Perm) midbody-stage cells showing that pAurB, pSC35, and SR domain proteins colocalize in ACBs (asynchronous cultures, 48 h after transfection with siNups or siControl co-stained for ACB components as indicated). (**B**) Immunofluorescence of nuclear speckles, MIGs, and ACBs in asynchronous conditions (as in **A**, but non-pre-permeabilized and at a variety of cell-cycle stages as designated). (**C**) Timeline of live-cell imaging of abscission timing following treatment with DMSO, 1 µm DYRK3i, or 1 µm CLK1i. (**D-E**) Immunofluorescence of midbody-stage cells fixed 60’ after addition of vehicle or inhibitors, with quantification of cytoplasmic foci marked by α-pSC35. N=300 midbodies/condition from n=3 biological replicates. **(F)** Timing from telophase to abscission (cells treated as in **C**). n=3 biological replicates.

Without the abscission checkpoint to induce transition to ACBs, newly discovered MIG factors, pAurB, pppCHMP4C, and CHMP4B relocalized individually to different sites in control cells at telophase. In the absence of abscission checkpoint activation, the checkpoint maintenance factors pAurB and pppCHMP4C relocalized into nuclei where, interestingly, they partially colocalized with nuclear speckles (Figure 4A, Figure 4—figure supplement 2A). In contrast, CHMP4B remained cytoplasmic and also concentrated at the midbody (Figure 3D, Figure 4— figure supplement 1B). Abscission checkpoint activation delayed timely nuclear relocalization of pAurB and pppCHMP4C, as well as the canonical MIG marker pSC35, until abscission or just prior (Figure 4A, Figure 4—figure supplement 2A). These observations indicate that individual components disperse to a variety of sites when MIGs/ACBs dismantle.

### ACBs regulate abscission timing

To probe for direct ACB functions in abscission delay, we tested whether abscission timing was altered when we artificially stimulated their formation/maintenance without otherwise triggering the abscission checkpoint. This was accomplished by inhibiting DYRK3, a kinase that prevents premature MIG formation in metaphase (Rai et al., 2018), or by inhibiting CLK1, a kinase that promotes component release from nuclear speckles and thus may similarly regulate MIG stability, allowing them to persist and at least partially mimic their maturation into ACBs (Araki et al., 2015; Colwill et al., 1996). CLK inhibition was previously reported to accelerate abscission (Petsalaki and Zachos, 2016), but our studies employed a CLK1/2-specific inhibitor rather than a pan-CLK inhibitor and our treatment windows were much shorter than those employed previously (30 minutes vs. 5 hours). Under these conditions, inhibition of either CLK1 or DYRK3 kinase increased the percentage of midbody cells with pSC35-positive foci in midbody-stage cells from 13% (control) to 23% (DYRK3i) or 21% (CLK1i) (Figure 4C-E), and increased median abscission times in the bulk population by 11 minutes (DYRK3i, 19% delay) or 6 minutes (CLK1i, 10% delay), as assayed in live cell imaging experiments (Figure 4F). Similarly, the time course of midbody resolution in a synchronized cell population was delayed upon treatment with either inhibitor (Figure 4—figure supplement 3). Thus, treatments that promote MIG/ACB formation and maintenance delay abscission in cells where the abscission checkpoint has not been otherwise activated. These observations indicate that ACBs have a functional role in abscission delay.

### ACB formation and ALIX recruitment delay occur in response to multiple mitotic errors and in different cell types

To confirm that ACBs are a broadly relevant abscission checkpoint mechanism, we investigated their formation in relation to other known methods of abscission checkpoint activation: replication stress, intercellular tension, and lagging chromatin bridges within the midbody (Lafaurie-Janvore et al., 2013; Mackay and Ullman, 2015; Steigemann et al., 2009). We observed that ACB levels significantly increased in midbody-stage cells when the abscission checkpoint was induced by replication stress (Figure 5—figure supplement 1) or intercellular tension (Figure 5—figure supplement 2), but not by chromatin bridges (Figure 5—figure supplement 3). Delayed ALIX midbody recruitment followed this same trend. Thus, ACB formation and delayed recruitment of ALIX to the midbody were correlated and were both seen under three out of four different conditions known to induce the abscission checkpoint.

To test whether this newly-identified checkpoint mechanism is deployed in other cellular contexts, we probed the non-transformed epithelial cell-line RPE1 for key checkpoint hallmarks.

RPE1 cells exhibited an abscission checkpoint in response to Nup depletion, as measured by an increase in midbody-stage cells and a delay in ALIX recruitment to the midbody (Figure 5— supplement 4A-C). This response was less robust than observed in HeLa cells, suggesting that corrective or compensatory pathways may abrogate errors that induce abscission arrest or that RPE1s may be less dependent on the ESCRT pathway. Nonetheless, ACBs were once again detected in nearly all midbody-stage RPE1 cells subjected to the checkpoint induction protocol, and contained pppCHMP4C, pAurB, CHMP4B, and pSC35 (Fig. 5A-C, Figure 5—figure supplement 4D). These observations demonstrate that checkpoint-dependent delay of ALIX midbody recruitment and ACB formation occur in distinct cell types and are broadly relevant as abscission checkpoint mechanisms.

**Figure 5.**
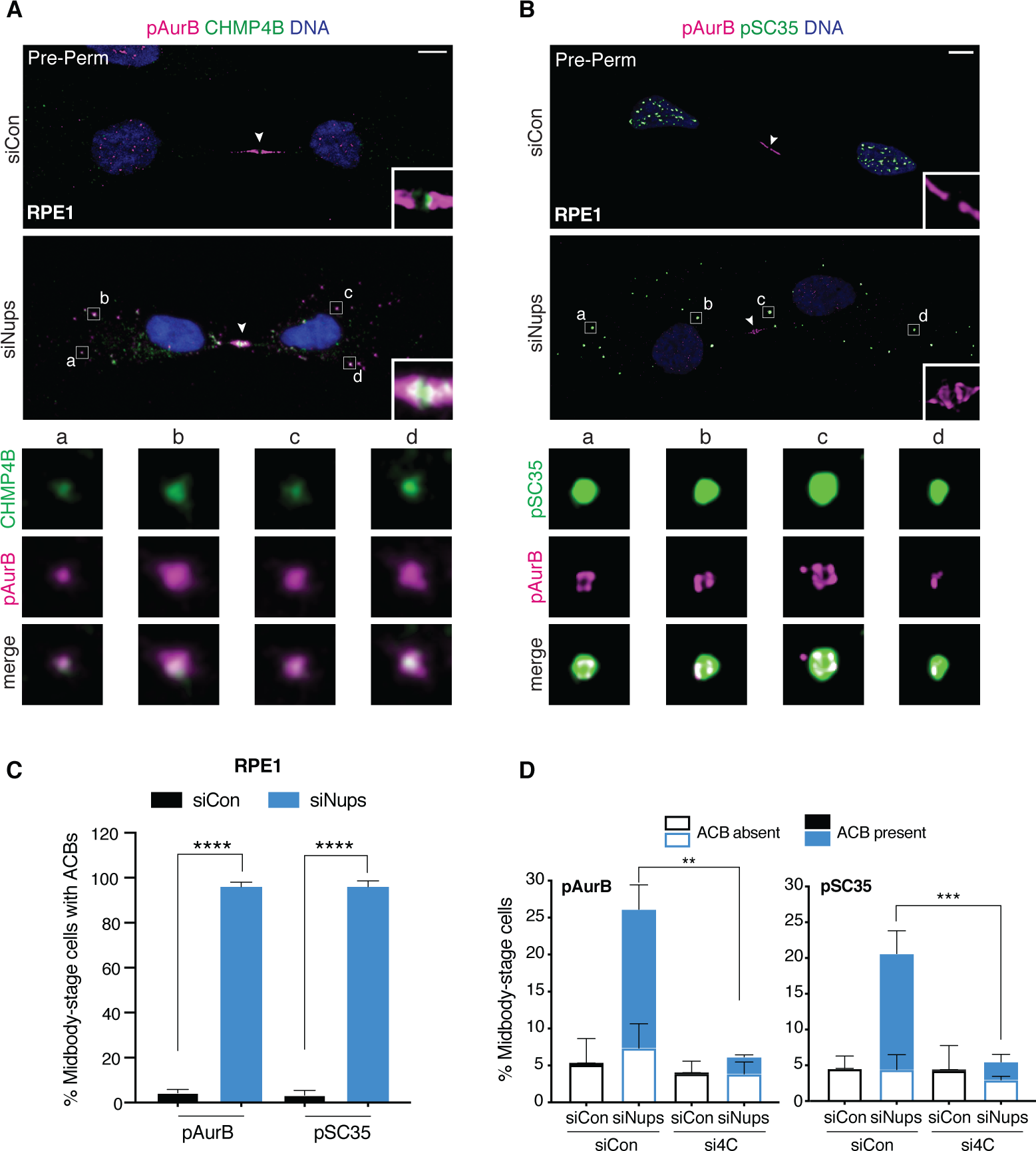
ACBs are conserved and dependent upon abscission checkpoint factor CHMP4C. (A-B) Confocal z-projections of pre-permeabilized control and checkpoint-activated RPE1 cells following 14 h thymidine release, stained for ACB markers as indicated. (C) Quantification of control and checkpoint activated RPE1 cells with ACBs 14 h post-thymidine release, detected by α-pAurB or α-pSC35. N=300 midbody-stage cells scored/condition from n=3 biological replicates. (D) Quantification of midbody-stage HeLa cells under asynchronous conditions (72 h after transfection with indicated siRNAs) containing ACB’s (marked by α-pAurB or α-pSC35). N= 300 midbody-stage cells/condition from n=3 biological replicates. P-values compare % midbodies with ACBs.

### ACB formation requires CHMP4C

To probe how the abscission checkpoint regulates ACBs, we depleted the key abscission checkpoint regulator CHMP4C and assayed ACBs. As expected, co-depletion of CHMP4C in Nup-depleted cells abrogated the checkpoint and reduced midbody-stage cells. This treatment also dramatically reduced the percentage of midbody-stage cells with ACBs (Figure 5D). The clear dependence on CHMP4C reinforces the connection between the abscission checkpoint and ACB appearance and suggests that ACB structures may depend on CHMP4C directly. The suppression of ACB formation in the absence of CHMP4C, despite depletion of Nups, also confirms that ACBs do not form due to altered function of the nuclear pore per se.

### ACBs contain ALIX and ALIX depletion reduces ACB size

To probe the connection between the presence of ACBs and the delay of ALIX recruitment to the midbody further, we tested whether ALIX itself might target to ACBs. To avoid the strong signal for ALIX present in the cytoplasm (Figure 2C), we again used pre-permeabilization conditions to remove soluble cytoplasmic content. Using this method, we were able to detect ALIX in ACBs in both HeLa and RPE1 cells (Figure 6A-B). This ACB localization was specific as ALIX knockdown decreased ALIX signal in ACBs (Figure 6—figure supplement 1A-B). Interestingly, although pAurB and pppCHMP4C clearly localize to MIGs (Figure 6—figure supplement 1E), we could not detect ALIX in these structures (Figure 6—figure supplement 1C-D). These observations indicate that ALIX is present in ACBs and, although we cannot rule out that our detection method is limiting for MIGs, it appears that ALIX is recruited to ACBs following their maturation from MIGs. In the absence of ALIX, ACB-like structures still persist in late midbody-stage cells in response to checkpoint activation, indicating that ALIX is not absolutely required for ACB formation (Figure 6C). However, ALIX depletion reduced the average ACB cross sectional area by ∼35% (corresponding to ∼50% reduction in volume), as measured using two different ACB markers (Figure 6D, Figure 6—figure supplement 2A). Depletion of ALIX did not translate into a marked reduction in total ACB area owing to a concomitant increase in the number of the focal ACB-like structures in checkpoint-activated midbody-stage cells (Figure 6—figure supplement 2B-C). Thus, ALIX is a component of ACBs and is required to create full-sized ACBs or to maintain their integrity.

**Figure 6.**
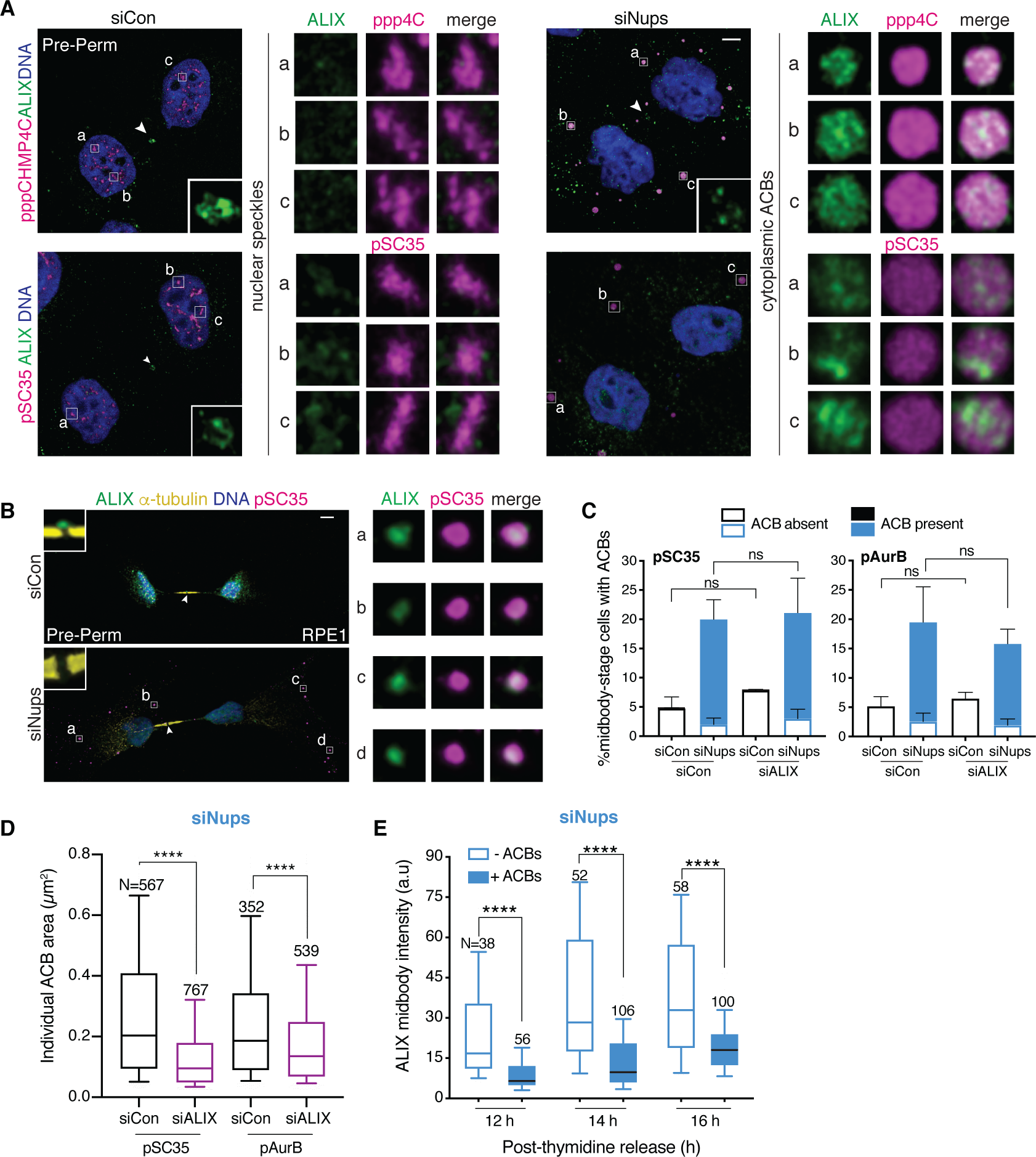
ACBs contain ALIX and cells with ACBs have delayed ALIX midbody recruitment. (**A**) Confocal z-projections of pre-permeabilized midbody-stage HeLa cells under asynchronous conditions (48 h after transfection with siNups or siControl), stained as indicated. (**B**) Confocal z-projections of pre-permeabilized control and checkpoint-activated RPE1 cells (14 h post-thymidine release) stained as indicated. (**C-D**) Quantification of (**C**) midbody-stage cells with/without ACBs detected with α-pAurB or α-pSC35, (**D**) ACB size under asynchronous conditions (72 h after transfection with indicated siRNAs). (**C**) N=300 midbodies scored/condition from n=3 biological replicates. P-values compare % midbodies with ACB’s. (**D)** n=3 biological replicates. (**E**) Time course quantification of relative ALIX midbody intensity in checkpoint-activated cells. Midbody-stage cells were binned into categories with or without ACB’s (marked by α-pAurB) using images from n=3 biological replicates.

### ACB formation correlates with reduced midbody ALIX levels

The presence of ALIX in ACBs suggested that the checkpoint-dependent delay in ALIX recruitment to the midbody may occur because a modified sub-population of ALIX that would normally be targeted to the midbody is instead retained in ACBs. To determine whether ACB formation and ALIX midbody recruitment delay are connected, cells were scored for both the presence of ACBs and the intensity of ALIX at the midbody, under conditions that synchronously activated the abscission checkpoint. Consistent with a direct ALIX sequestration model, we found that the presence of ACBs strongly correlated with decreased ALIX levels at the midbody in individual cells (Fig. 6E, Figure 6—figure supplement 1D). This correlation was striking even late in the post-thymidine time course, when overall ALIX midbody recruitment has begun to recover.

## Discussion

We have developed a new abscission checkpoint synchronization assay ideally suited to elucidating the regulation of abscission timing. Employing this assay, we found that checkpoint activation delays ALIX recruitment to the midbody and identified Abscission Checkpoint Bodies (ACBs) as a previously uncharacterized cytoplasmic body that contributes a new facet of abscission regulation. Specifically, we find that the abscission checkpoint restrains recruitment of ALIX and the downstream factor IST1 to the midbody, helping to explain how abscission is stalled in response to mitotic errors. Furthermore, we identified ACBs as a checkpoint-dependent compartment that concentrates splicing, checkpoint and abscission factors, including pSC35, pAurB, pppCHMP4C, CHMP4B, and ALIX. ACBs appear to be derived from MIGs, a compartment whose components and cell cycle behavior had previously been partially characterized, but whose functional role(s) are unknown at a mechanistic level (Ferreira et al., 1994; Prasanth et al., 2003; Rai et al., 2018; Reuter et al., 1985; Spector and Lamond, 2011; Tripathi and Parnaik, 2008; Turner and Franchi, 1987). Although much remains to be learned about MIG/ACB architecture and function, ACB formation/maintenance requires the abscission checkpoint factor CHMP4C, with a distinct role for the cytokinesis factor ALIX in their maturation. CHMP4C and pAurB colocalize within ACBs, which may favor formation and maintenance of the pppCHMP4C isoform because CHMP4C is an AurB substrate (Capalbo et al., 2012; Carlton et al., 2012). At least two ACB components, CHMP4C and CHMP4B, can bind ALIX (McCullough et al., 2008), and CHMP4C mutations that cripple or eliminate ALIX binding impair the checkpoint (Sadler et al., 2018), highlighting the functional importance of the CHMP4C-ALIX interaction in checkpoint maintenance, and raising the possibility that their interaction could target either the CHMP4 proteins or ALIX to ACBs.

In functional terms, targeting of ALIX to ACBs may sequester a particular subpopulation of this protein away from the midbody, contributing to the checkpoint-dependent abscission delay. Such targeting is predicted to be halted when the checkpoint is satisfied and phosphatases such as PP1 counteract pAurB and thus pppCHMP4C (Bhowmick et al., 2019; Capalbo et al., 2019). Previous studies have demonstrated that ALIX must be activated by phosphorylation on its autoinhibitory C-terminal tail in order to bind CHMP4 proteins and to function in abscission (Sun et al., 2016; Zhai et al., 2011). pALIX is therefore an attractive candidate for the subpopulation of ALIX that is sequestered within ACBs.

The key role of ALIX in the abscission of cultured, transformed cells is now well established (Carlton and Martin-Serrano, 2007; Morita et al., 2007), and is reinforced by recent studies elucidating its stepwise recruitment to the abscission zone, where it works in a complex with syndecan4 and syntenin to recruit other ESCRT factors required for the scission event (Addi et al., 2020). However, newly reported mouse knockout models reveal a more complex picture in vivo. In these mouse models, ALIX and CEP55 were found to be required for normal brain and kidney development and cell division, whereas other organs appear largely unaffected (Campos et al., 2016; Tedeschi et al., 2020). Moreover, humans without functional ALIX display microcephaly and kidney defects but are otherwise healthy and can live into their 20s (Khan et al., 2020). We reconcile these observations with our model of the abscission checkpoint in the following ways: 1) Redundant pathways may ensure that abscission (and an abscission checkpoint) takes place in vivo; indeed, a recent report argues that ESCRT proteins, including ALIX, are still recruited to the midbody in the CEP55 knockout mouse (Little et al., 2020), and in cases where ALIX is absent, TSG101 may dominate in driving ESCRT activity (Christ et al., 2016; Karasmanis et al., 2019). 2) It is also possible that tumor-derived cells are more dependent on the ESCRT pathway for abscission than non-cancerous cells, albeit with the exception of the developing brain and kidney. In line with this, ESCRT factors are often overexpressed in cancer (Jeffery et al., 2015; Lin et al., 2020), and our results show a more robust abscission checkpoint response in HeLa cells compared to non-transformed RPE1 cells. This important issue requires further study, but could potentially be used to advantage in therapeutic approaches.

ALIX targeting to ACBs is the first known cytoplasmic mechanism for abscission checkpoint regulation, but several lines of evidence indicate that this is just one of a repertoire of mechanisms that can be integrated to accomplish abscission delay. For example, the ACB mechanism does not function measurably when the checkpoint is induced by lagging chromatin bridges (Figure 5—figure supplement 3). Consistent with that observation, ALIX does not appear delayed in its recruitment to the midbody under these circumstances. Thus, other sensing and signaling mechanisms must be responsible for the AurB-dependent abscission delay triggered by chromosomal bridges. Indeed, a histone acetyltransferase complex plays an integral role in response to chromatin in the cell cleavage plane in the analogous NoCut checkpoint in yeast (Mendoza et al., 2009). There is also other precedent for modularity in abscission checkpoint mechanisms elicited by different cues. For example, although phosphorylation of IST1 by ULK3 is required for an abscission delay in response to chromatin bridges or Nup depletion, it does not appear to be required for tension-mediated abscission regulation (Caballe et al., 2015). Furthermore, other abscission delay mechanisms, such as ANCHR-dependent sequestration of the ATPase VPS4 away from the abscission zone (Thoresen et al., 2014), may be deployed in particular combinations depending on the error present.

Finally, our demonstration that late midbody-stage ACBs are derived from telophase MIGs begs the question of why these bodies colocalize factors that function in both abscission and mRNA biogenesis. Interphase nuclear speckles are hypothesized to be transcription hubs that mediate efficient splicing especially of high-copy genes, and molecular detail of their function is emerging (Smith et al., 2020). The functional roles of MIGs are currently less clear. Abnormal MIG assembly triggers metaphase arrest (Rai et al., 2018; Sharma et al., 2010), implying a role in mitotic progression, but the mechanism is not yet understood. We have shown that artificially promoting ACB assembly triggers an abscission delay (Figure 4D-F, Figure 4—figure supplement 3), indicating that ACBs have a role in cytokinetic progression. Intriguingly, our observations suggest that ACBs are normally resolved and nuclear speckles are reformed before abscission takes place. It will therefore be of interest to determine whether sequestration and coordinated release of the splicing factors or other regulatory factors present within ACBs are required for abscission regulation and cytokinetic progression.

## Materials and Methods

### Antibodies

Details regarding antibodies used in this study can be found in Supplementary Table 1.

### Cell Culture

HeLa cells were cultured and maintained at 37°C and 5% CO_2_ in DMEM supplemented with 10% FBS. RPE1 cells were supplemented with 10 µg/ml hygromycin (Invitrogen, Carlsbad, CA) to maintain hTERT expression. The HA-CHMP4C HeLa cell line (Carlton et al., 2012) was supplemented with 100 µg/ml G418 (ThermoFisher, Waltham, MA). At the outset of these studies, HeLa and RPE1 cells were tested and found negative for mycoplasma using a PCR mycoplasma detection kit (ABM, Bellingham, WA). Cell types were authenticated by sequencing 24 loci (University of Utah Sequencing Core).

### siRNA Transfections

Cells were singly transfected with siRNA for 42-72 h, as indicated in figure legends, using Lipofectamine RNAiMax (ThermoFisher) according to the manufacturer’s instructions. Media was exchanged 24 h after transfection and cells were incubated between 18-48 h until harvesting. For co-transfections, cells were seeded with combinations of 1) siALIX, siCon, and siNup153/50 or 2) siCHMP4C, siCon, and siNup153/50. Media was exchanged 24 h after transfection. Cells were fixed 72 h post-transfection to allow adequate time for CHMP4C or ALIX knockdown. Details regarding siRNAs used in this study are provided in Supplementary Table 2.

### Immunofluorescence

Cells were seeded on acid-washed, 10 µg/ml fibronectin-treated, glass coverslips and then treated according to the individual experimental protocol. To fix, cells were removed from the incubator, washed once in PBS pH 7.4, and then fixed by one of three methods as notated in Table S1:

1. 10 min, −20°C methanol,
2. 15 minutes, 4°C, 2-4% paraformaldehyde (PFA) followed by 5 minutes, 23°C, 0.5% Triton-X in PBS
3. rinsed with PHEM buffer at 23°C (50 mM PIPES, 25 mM HEPES pH 7.0, 10 mM EGTA, 4 mM MgSO_4_, with PMSF (1 mM), aprotinin, leupeptin added freshly), pre-permeabilized with 0.5% Triton-X in PBS for 1 minute at 23°C, fixed with 2-4% ice cold PFA for 15 minutes, and incubated with 0.5% Triton-X in PBS for 5 minutes at 23°C.

After fixing, cells were rinsed twice with 2 ml PBS each time and then blocked (3% FBS and 0.1% Triton X-100 in PBS) for 30 min on the bench top. Primary antibodies were applied at the dilution noted in Table S1 for at least 1 h, 23°C in blocking solution. After 1 wash with 2 ml PBS, secondary antibodies (Thermofisher) were applied for 45 min-1 h, and cells were washed in 2 ml PBS. For Figure 2E and Figure 2—figure supplement 2C, DNA was stained with Hoechst for 10 minutes at 23°C. For all other images, coverslips were mounted with ProLong Gold Antifade Reagent with DAPI (Thermofisher) on a microscope slide.

### Imaging

Images were acquired using three different microscopes:

1. Leica SP8 Confocal 63x 1.4 oil HC PL APO objective with adjustable white-light laser to control for bleed-through (Figure 3D, Figure 4A, Figure 5B-C, Figure 6A-B, Figure 3—figure supplement 2A, Figure 4—figure supplement 1C, Figure 4—figure supplement 2A, Figure 6— figure supplement 1A, C, Figure 6—figure supplement 2A). Images were acquired as z-stacks and each individual slice was deconvolved using the Hyvolution and Lightening modes on Leica App Suite X Software. Presented images are maximum z-projections of the deconvolved slices.
2. Zeiss Axioskop 2 Widefield 63x 1.4 oil DIC Plan Apochromat objective (Figure 1E, Figure 2A, C, Figure 3A, Figure 4B, Figure 2—figure supplement 2A, Figure 2—figure supplement 3A, Figure 3—figure supplement 3B, E, G, Figure 4—figure supplement 1A-B, Figure 5—figure supplement 1C, E, Figure 5—figure supplement 2C, E, Figure 5—figure supplement 4D, and Figure 6—figure supplement 1E-F). Images were acquired as single plane widefield images.
3. Axioskop 2 mot PLUS, 63x 1.4 oil and 100x oil Plan-Apochromat objectives (Figure 2E, Figure 4D, Figure 2—figure supplement 2C, Figure 4—figure supplement 3E, and Figure 5— figure supplement 3A, C). These images were acquired as single plane widefield images.

### Cell and Image Scoring

Midbody-stage cells were identified by α-tubulin staining and when counting cell populations were always counted as one cell. Cell populations were counted and sorted into interphase, midbody-stage, mitotic, recently abscised pair (counted as one cell), multinucleate, and failed bridge (counted as one cell). Cells were counted unblinded while on the microscope.

Midbodies were identified as “early” or “not early” using α-tubulin staining before scoring for the examined phenotype. Midbodies were classified as early based on midbody width, level of midbody-pinching, nuclear area, and flatness of cells, see Figure 4B for illustration.

Midbodies were scored as having ACBs if they had at least one ACB. These cases were rare, however, because most midbody-stage cells with ACBs, had 10-40 ACBs. For most figures, cells were counted as having or not having ACBs, unblinded while on the microscope. In Figure 3—figure supplement 2B, ACBs were counted from images using “Find Maxima” with Fiji software (NIH). To quantify ALIX signal in ACBs: deconvolved z-slices were stacked and then uniformly adjusted for brightness and contrast. The pSC35 or pAurB channels were uniformly thresholded, and then cytoplasmic objects between 0.025-5 µm^2^ were marked as Regions Of Interest (ROIs). ROIs were used to measure individual ACB intensity and area in the non-thresholded ALIX channel. Values were background corrected with large ROIs in the cytoplasm that excluded ACBs.

Quantification of fluorescence staining intensity at the midbody was also done with Fiji software (NIH). The freehand selection tool was used to outline the region of interest at the midbody, and staining intensity within this area was measured. Signals were background corrected using measurements from adjacent regions. In cases where the protein was not visible at the midbody, these methods sometimes generated a small negative intensity value after background correction because the Flemming body is a natural dark zone. In these cases, intensity was valued at zero.

### Immunoblotting

Cells were lysed in NP40 lysis buffer (50 mM Tris pH 7.4, 250 mM NaCl, 5 mM EDTA, 50 mM NaF, 1 mM Na_2_VO_4_, 1% Nonidet P40, 0.2% NaN_3_, with freshly added PMSF (1 mM), aprotinin, leupeptin) for 30 minutes, 4°C, vortexing every 10 minutes. Lysates were clarified by spinning at 13,000 g for 10 minutes at 4°C and protein contents in clarified lysates were quantified by Bradford assay before gel loading. Nup153 was detected on a 5% SDS gel and both HSP90 and Nup50 were detected on a 12% SDS gel. 30 µg lysate per sample were prepared with SDS loading buffer, resolved by SDS-PAGE, and wet-transferred to PVDF membranes for 2 hours at 40 V. Membranes were blocked with 5% milk in Tris-buffered saline (TBS) for at least 30 minutes, 23°C, and were incubated with primary antibodies in 5% milk in TBS-T (0.1% Tween 20/TBS) overnight at 4°C. Membranes were washed in TBS-T and incubated with the corresponding secondary antibodies conjugated with HRP (Thermofisher) in TBS-T for 1 h, 23°C, and washed again. Blots were detected with Western Lightning PLUS ECL (PerkinElmer, Waltham, MA) on Hyblot CL film (Thomas Scientific, Swedesboro, NJ).

### Checkpoint Activation

The abscission checkpoint was activated using one of four different methods.

1. Nup Depletion: cells were seeded on glass coverslips and transfected with 10 nM siNup153 and 10 nM siNup50 or control siRNA. In asynchronous experiments, media was exchanged at 24 h, and cells were fixed at 48 h or 72 h. In synchronous experiments, 2 mM thymidine (Calbiochem, San Diego, CA) was added to samples 8 hours post-seeding. Cells were incubated with thymidine for 24 h and then washed with PBS 3 times to remove thymidine and siRNA transfection mixture. Fresh media was added, and cells were harvested 10-18 h later. In cases where cells were treated with AurB inhibitor (AurBi), 2 µm ZM 447439 (Ditchfield et al., 2003)(Biotechne, Minneapolis, MN) or DMSO vehicle was added 30 minutes (for partial abscission completion in Fig. 2G, Figure 2—figure supplement 3C) or 1 hour (for almost complete abscission completion in Figure 1D-E) prior to fixing.
2. Chromatin Bridges: cells were seeded on acid-washed glass coverslips and incubated for 48 h without perturbation. Cells were fixed and stained with Lap2ß to identify the naturally-occurring small percentage of cells with lagging chromatin. Phenotypes were scored in cells with and without lagging chromatin.
3. Replication Stress: following the general procedure of Mackay and Ullman, 2015, cells were seeded on glass coverslips, and 2 mM thymidine was added 8 h post-seeding to synchronize cells. Cells were incubated for 24 h, and cells were washed 3 times with 2ml PBS to remove thymidine. Fresh media was added and supplemented with 0.4 µM aphidicolin or DMSO. Cells were harvested 12-18 hours later.
4. Tension: following the general procedure of Lafaurie-Janvore et al., 2013, in which intercellular tension is limited by cell density, cells were seeded on glass coverslips at a density of 25,000 cells/ml (low density, high tension) or 75,000 cells/ml (high density, low tension). Cells were harvested at 48 h post-seeding without additional perturbations. Cells were only scored as “high tension” if at least one side was not touching a neighbor and was free to expand and scored as “low tension” if completely surrounded by other cells and not free to spread.

### ACB Induction Experiments

For fixed-imaging time courses, HeLa cells were seeded on glass coverslips in 24-well dishes, and 2 mM thymidine was added 8 hours post-seeding. After 24 h, thymidine was washed out with PBS. 13 hours post-thymidine release, after most cells had completed metaphase, 1 µM DYRK3 inhibitor (GSK 626616, Tocris, Minneapolis, MN), 1 µM CLK1 inhibitor (534350, Millipore, Burlington, MA), or DMSO were each added individually to 6 separate wells.

Coverslips from wells corresponding to each treatment were fixed directly after inhibitor/vehicle was added for the first timepoint at 13 h post-thymidine release and then at hourly intervals until 18 h post-thymidine release.

Live-imaging experiments used HeLa cells expressing H2B-mCherry and GFP-α-tubulin. Cells were seeded in a Lab-Tek II 8-chambered #1.5 German Coverglass System and incubated for 48 hours. Stage positions were first set on the microscope, and then 1 µM DYRK3i, 1 µM CLK1i, or DMSO were added just prior to initiation of imaging to monitor abscission timing.

Imaging was carried out for 3 h on a Nikon Ti-E widefield inverted microscope (Nikon 20x N.A. dry objective lens) equipped with Perfect Focus system and housed in a 37°C chamber (OKOLAB, Ambridge, PA) with 5% CO_2_. Multiple fields of view were selected at various x and y coordinates, and images were acquired using a high sensitivity Andor Zyla CMOS camera (Andor, Manchester, CT) controlled by NIS-Elements software. Images were acquired every 5 min, and abscission time was measured as the time from midbody formation to disappearance. In parallel, cells were seeded on glass coverslips, incubated for 48 hours, and either DYRK3i, CLK1i, or DMSO were added for 60 minutes prior to fixing and detection of pSC35 or pAurB foci.

### Statistical Analysis

Four statistical tests were used to evaluate significance of data. When comparing two samples, p-values were calculated using an unpaired t-test (Fig. 6E, Figure 5—figure supplement 3A, B, D, Figure 5—figure supplement 3D, Figure 5—figure supplement 4A, Figure 6—figure supplement 1B, D). When comparing two complete time curves, p-values were calculated using Welch’s t-test (Figure 3C). When comparing datasets with three or more samples, p-values were calculated using one-way ANOVA with Sidak’s multiple comparisons test (Figure 2F-G, Figure 3E, Figure 4E-F, Figure 5A, D, Figure 6C-E, Figure 2—figure supplement 1C, Figure 2—figure supplement 2D, Figure 2—figure supplement 3C, Figure 3—figure supplement 2B, Figure 4—figure supplement 3C-D, Figure 5—figure supplement 3B, Figure 6—figure supplement 2B-C). When comparing two or more datasets over a time course or multiple categories within a sample, p-values were calculated using two-way ANOVA with Sidak’s multiple comparisons test (Figure 3—figure supplement 1G, Figure 4—figure supplement 3B Figure 5—figure supplement 1B, D, F, Figure 5—figure supplement 4C).

## Acknowledgments

We thank F. Barr and P. D’Avino for antibodies, and J. Martin-Serrano for the HA-CHMP4C cell line. We thank M. Smith and D. Wenzel for expert advice on microscopy and biochemistry, respectively. Leica Confocal SP8 and Nikon Automated Widefield microscopes were provided by the University of Utah Cell Imaging Core. We acknowledge funding from NIH R01GM112080 (K.S.U., W.I.S.), as well as financial support provided by the Huntsman Cancer Foundation and the Cell Response and Regulation Program at Huntsman Cancer Institute. We also acknowledge support by the National Cancer Institute of the National Institutes of Health under Award Number P30CA042014 for shared resources.

## Supplementary Materials

Figures 1S1-6S3

Tables S1-S2

## References

Addi C, Presle A, Frémont S, Cuvelier F, Rocancourt M, Milin F, Schmutz S, Chamot-Rooke J, Douché T, Duchateau M, Gianetto QG, Ménager H, Matondo M, Zimmermann P, Gupta-Rossi N, Echard A. 2020. The Flemmingsome reveals an ESCRT-to-membrane coupling required for completion of cytokinesis. Nat Commun 11. doi:10.1101/2020.01.15.907857

Agromayor M, Carlton J, Phelan J, Matthews D, Carlin L, Ameer-Beg S, Bowers K, Martin-Serrano J. 2009. Essential role of hIST1 in cytokinesis. MBoC 20:1374–1387. doi:10.1091/mbc.E08

Araki S, Dairiki R, Nakayama Y, Murai A, Miyashita R, Iwatani M, Nomura T, Nakanishi O. 2015. Inhibitors of CLK protein kinases suppress cell growth and induce apoptosis by modulating pre-mRNA splicing. PLoS One 10:1–18. doi:10.1371/journal.pone.0116929

Bajorek M, Morita E, Skalicky JJ, Morham SG, Babst M, Sundquist WI. 2009a. Biochemical analyses of human IST1 and its function in cytokinesis. MBoC 20:1360–1373. doi:10.1091/mbc.E08

Bajorek M, Schubert HL, McCullough J, Langelier C, Eckert DM, Stubblefield W-MB, Uter N, Myszka D, Hill C, Sundquist W. 2009b. Structural basis for ESCRT-III protein autoinhibition. Nat Struct Mol Biol 16:754–62. doi:10.1038/nsmb.1621

Bastos R, Barr F. Plk1 negatively regulates Cep55 recruitment to the midbody to ensure orderly abscission. JCB 191:751–760. doi:10.1083/jcb.201008108

Beck JS. 1961. Variations in the morphological patterns of “autoimmune” nuclear fluorescence. Lancet 277:1203–1205. doi:10.1016/S0140-6736(61)91944-4

Bhowmick R, Thakur RS, Venegas AB, Liu Y, Nilsson J, Barisic M, Hickson ID. 2019. The RIF1-PP1 Axis Controls Abscission Timing in Human Cells. Curr Biol 29:1232-1242.e5. doi:10.1016/j.cub.2019.02.037

Caballe A, Wenzel D, Agromayor M, Alam S, Skalicky J, Kloc M, Carlton J, Labrador L, Sundquist W, Martin-Serrano J. 2015. ULK3 regulates cytokinetic abscission by phosphorylating ESCRT-III proteins. Elife 4:1–36. doi:10.7554/eLife.06547

Campos Y, Qiu X, Gomero E, Wakefield R, Horner L, Brutkowski W, Han YG, Solecki D, Frase S, Bongiovanni A, D’Azzo A. 2016. Alix-mediated assembly of the actomyosin-tight junction polarity complex preserves epithelial polarity and epithelial barrier. Nat Commun 7:11876. doi:10.1038/ncomms11876

Capalbo L, Bassi ZI, Geymonat M, Todesca S, Copoiu L, Enright AJ, Callaini G, Riparbelli MG, Yu L, Choudhary JS, Ferrero E, Wheatley S, Douglas ME, Mishima M, D’Avino PP. 2019. The midbody interactome reveals unexpected roles for PP1 phosphatases in cytokinesis. Nat Commun 10:1–17. doi:10.1038/s41467-019-12507-9

Capalbo L, Mela I, Abad MA, Jeyaprakash AA, Edwardson JM, Avino PPD. 2016. Coordinated regulation of the ESCRT-III component CHMP4C by the chromosomal passenger complex and centralspindlin during cytokinesis. Open Biol 6:160248.

Capalbo L, Montembault E, Takeda T, Bassi ZI, Glover DM, D’Avino PP. 2012. The chromosomal passenger complex controls the function of endosomal sorting complex required for transport-III Snf7 proteins during cytokinesis. Open Biol 2:120070. doi:10.1098/rsob.120070

Carlton J, Agromayor M, Martin-Serrano J. 2008. Differential requirements for Alix and ESCRT-III in cytokinesis and HIV-1 release. PNAS 105:10541–10546. doi:10.1073/pnas.0802008105

Carlton J, Caballe A, Agromayor M, Kloc M, Martin-Serrano J. 2012. ESCRT-III governs the Aurora B-mediated abscission checkpoint through CHMP4C. Science 336:220–225. doi:10.1126/science.1217180.ESCRT-III

Carlton J, Martin-Serrano J. 2007. Parallels between cytokinesis and retroviral budding: a role for the ESCRT machinery. Science 316:1908–1912. doi:10.1126/science.1143422

Chen Y, Belmont AS. 2019. Genome organization around nuclear speckles. Curr Opin Genet Dev 55:91–9. doi:10.1016/j.gde.2019.06.008

Christ L, Wenzel EM, Liestøl K, Raiborg C, Campsteijn C, Stenmark H. 2016. ALIX and ESCRT-I/II function as parallel ESCRT-III recruiters in cytokinetic abscission. JCB 212:499–513. doi:10.1083/jcb.201507009

Colwill K, Pawson T, Andrews B, Prasad J, Manley J, Bell J, Duncan P. 1996. The Clk/Sty protein kinase phosphorylates SR splicing factors and regulates their intranuclear distribution. EMBO J 15:265–275.

Ditchfield C, Johnson VL, Tighe A, Ellston R, Haworth C, Johnson T, Mortlock A, Keen N, Taylor SS. 2003. Aurora B couples chromosome alignment with anaphase by targeting BubR1, Mad2, and Cenp-E to kinetochores. JCB 161:267–80. doi:10.1083/jcb.200208091

Ferreira JA, Carmo-Fonseca M, Lamond AI. 1994. Differential interaction of splicing snRNPs with coiled bodies and interchromatin granules during mitosis and assembly of daughter cell nuclei. JCB 126:11–23. doi:10.1083/jcb.126.1.11

Fu XD, Maniatis T. 1990. Factor required for mammalian spliceosome assembly is localized to discrete regions in the nucleus. Nature 343:437–441. doi:10.1038/343437a0

Gatta AT, Carlton JG. 2019. The ESCRT-machinery: closing holes and expanding roles. Curr Opin Cell Biol 59:121–132. doi:10.1016/j.ceb.2019.04.005

Hartwell L, Weinert T. 1989. Checkpoints : controls that ensure the order of cell cycle events. Science 246:629–34.

Hegemann B, Hutchins JRA, Hudecz O, Novatchkova M, Rameseder J, Sykora MM, Liu S, Mazanek M, Lenart P, Hériché J, Poser I, Kraut N, Hyman AA, Yaffe MB, Peters J-M. 2014. Systematic phosphorylation analysis of human mitotic protein complexes. Sci Signal 4. doi:10.1126/scisignal.2001993.Systematic

Jeffery J, Sinha D, Srihari S, Kalimutho M, Khanna KK. 2015. Beyond cytokinesis: the emerging roles of CEP55 in tumorigenesis. Oncogene 1–8. doi:10.1038/onc.2015.128

Karasmanis EP, Hwang D, Nakos K, Bowen JR, Angelis D, Spiliotis ET. 2019. A septin double ring controls the spatiotemporal organization of the ESCRT machinery in cytokinetic abscission. Curr Biol 29:1–9. doi:10.2139/ssrn.3291327

Kettenbach AN, Schweppe DK, Faherty BK, Pechenick D, Pletnev A a, Gerber S a. 2011. Quantitative phosphoproteomics identifies substrates and functional modules of Aurora and polo-like kinase activities in mitotic cells. Sci Signal 4:rs5. doi:10.1126/scisignal.2001497

Khan A, Alaamery M, Massadeh S, Obaid A, Kashgari AA, Walsh, Christopher A, Eyaid W. 2020. PDCD6IP, encoding a regulator of the ESCRT complex, is mutated in microcephaly. Clin Genet. doi:10.1111/1744-1633.12020

Lafaurie-Janvore J, Maiuri P, Wang I, Pinot M, Manneville J-B, Betz T, Balland M, Piel M. 2013. ESCRT-III assembly and cytokinetic abscission are induced by tension release in the intercellular bridge. Science 339:1625–1629. doi:10.1126/science.1233866

Lee H, Elia N, Ghirlando R, Lippincott-Schwartz J, Hurley J. 2008. Midbody Targeting of the ESCRT Machinery by a Noncanonical Coiled Coil in CEP55. Science 322:576–580. doi:10.1126/science.1162042

Li H, Bingham PM. 1991. Arginine/serine-rich domains of the su(wa) and tra RNA processing regulators target proteins to a subnuclear compartment implicated in splicing. Cell 67:335– 342. doi:10.1016/0092-8674(91)90185-2

Lin S, Wang M, Cao Q, Li Q. 2020. Chromatin modified protein 4C (CHMP4C) facilitates the malignant development of cervical cancer cells. FEBS Open Bio. doi:10.1002/2211-5463.12880

Little JN, McNeely KC, Michel N, Bott CJ, Lettieri KS, Hecht MR, Martin SA, Dwyer ND, Janosik SM. 2020. Loss of coiled-coil protein CEP55 impairs abscission processes and results in p53-dependent apoptosis in devleoping cortex. bioRxiv 42:1. doi:10.1017/CBO9781107415324.004

Maciejowski J, Li Y, Bosco N, Campbell PJ, Maciejowski J, Li Y, Bosco N, Campbell PJ, Lange T De. 2015. Chromothripsis and Kataegis Induced by Telomere Article Chromothripsis and Kataegis Induced by Telomere Crisis. Cell 163:1641–1654. doi:10.1016/j.cell.2015.11.054

Mackay D, Makise M, Ullman K. 2010. Defects in nuclear pore assembly lead to activation of an Aurora B-mediated abscission checkpoint. JCB 191:923–931. doi:10.1083/jcb.201007124

Mackay D, Ullman K. 2015. ATR and a Chk1-Aurora B pathway coordinate postmitotic genome surveillance with cytokinetic abscission. MBoC 26:2217–2226. doi:10.1091/mbc.E14-11-1563

McCullough J, Fisher R, Whitby F, Sundquist W, Hill C. 2008. ALIX-CHMP4 interactions in the human ESCRT pathway. PNAS 105:7687–7691. doi:10.1073/pnas.0801567105

Mendoza M, Norden C, Durrer K, Rauter H, Uhlmann F, Barral Y. 2009. A mechanism for chromosome segregation sensing by the NoCut checkpoint. Nat Cell Biol 11:477–483. doi:10.1038/ncb1855

Mintz PJ, Patterson SD, Neuwald AF, Spahr CS, Spector DL. 1999. Purification and biochemical characterization of interchromatin granule clusters. EMBO J 18:4308–4320. doi:10.1093/emboj/18.15.4308

Morita E, Sandrin V, Chung H-Y, Morham S, Gygi S, Rodesch C, Sundquist W. 2007. Human ESCRT and ALIX proteins interact with proteins of the midbody and function in cytokinesis. EMBO J 26:4215–4227. doi:10.1038/sj.emboj.7601850

Norden C, Mendoza M, Dobbelaere J, Kotwaliwale C V., Biggins S, Barral Y. 2006. The NoCut pathway links completion of cytokinesis to spindle midzone function to prevent chromosome breakage. Cell 125:85–98. doi:10.1016/j.cell.2006.01.045

Petsalaki E, Zachos G. 2019. Building bridges between chromosomes: novel insights into the abscission checkpoint. Cell Mol Life Sci 76:4291–4307. doi:10.1007/s00018-019-03224-z

Petsalaki E, Zachos G. 2016. Clks 1, 2 and 4 prevent chromatin breakage by regulating the Aurora B-dependent abscission checkpoint. Nat Commun 7:11451. doi:10.1038/ncomms11451

Pharoah P, Tsai Y, Ramus S, Phelan C, Ellen L, Lawrenson K, Price M, Fridley B, Tyrer J, Weber R, Karevan R, Larson M, Song H, Daniel C, Bacot F, Vincent D, Cunningham J, Dennis J. 2013. GWAS meta-analysis and replication identifies three new susceptibility loci for ovarian cancer. Nat Genet 45:362–370. doi:10.1038/ng.2564.GWAS

Prasanth K V., Sacco-Bubulya PA, Prasanth SG, Spector DL. 2003. Sequential entry of components of gene expression machinery into daughter nuclei. MBoC 14:1043–1057. doi:10.1091/mbc.E02

Rai AK, Chen J-X, Selbach M, Pelkmans L. 2018. Kinase-controlled phase transition of membraneless organelles in mitosis. Nature 559:211–216. doi:10.1038/s41586-018-0279-8

Reuter R, Appel B, Rinke J, Luhrmann R. 1985. Localization and structure of snRNPs during mitosis: immunofluorescent and biochemical studies. Exp Cell Res 159:63–79.

Sadler JBA, Wenzel DM, Williams LK, Guindo-mart M, Alam SL, Maria J, Torrents D, Ullman KS, Sundquist WI, Martin-serrano J. 2018. A cancer-associated polymorphism in ESCRT-III disrupts the abscission checkpoint and promotes genome instability. PNAS 115:E8900–8908.

Saitoh N, Sakamoto C, Hagiwara M, Agredano-Moreno LT, Jimenez-Garcia LF, Nakao M. 2012. The distribution of phosphorylated SR proteins and alternative splicing are regulated by RANBP2. MBoC 23:1115–1128. doi:10.1091/mbc.E11-09-0783

Saitoh N, Spahr CS, Patterson SD, Bubulya P, Neuwald AF, Spector DL. 2004. Proteomic analysis of interchromatin granule clusters. MBoC 15:3876–3890. doi:10.1091/mbc.E04

Sharma A, Takata H, Shibahara K, Bubulya A, Bubulya PA. 2010. Son is essential for nuclear speckle organization and cell cycle progression. MBoC 21:650–663. doi:10.1091/mbc.E09

Skop A, Liu H, Yates J, Meyer B, Heald R. 2004. Dissection of the mammalian midbody proteome reveals conserved cytokinesis mechanisms. Science 305:61–66.

Smith KP, Hall LL, Lawrence JB. 2020. Nuclear hubs built on RNAs and clustered organization of the genome. Curr Opin Cell Biol 64:67–76. doi:10.1016/j.ceb.2020.02.015

Spector D, Lamond A. 2011. Nuclear Speckles. Cold Spring Harb Perspect Biol 3:a000646. doi:10.1007/s11748-010-0709-5

Spector DL, Smith HC. 1986. Redistribution of U-snRNPs during mitosis. Exp Cell Res 163:87–94.

Steigemann P, Wurzenberger C, Schmitz MH a, Held M, Guizetti J, Maar S, Gerlich DW. 2009. Aurora B-mediated abscission checkpoint protects against tetraploidization. Cell 136:473–484. doi:10.1016/j.cell.2008.12.020

Strom AR, Brangwynne CP. 2019. The liquid nucleome – phase transitions in the nucleus at a glance. JCS 132:jcs235093. doi:10.1242/jcs.235093

Sun S, Sun L, Zhou X, Wu C, Wang R, Lin S-H, Kuang J. 2016. Phosphorylation-dependent activation of the ESCRT function of ALIX in cytokinetic abscission and retroviral budding. Dev Cell 36:331–343. doi:10.1016/j.devcel.2016.01.001

Tedeschi A, Almagro J, Renshaw MJ, Messal HA, Behrens A, Petronczki M. 2020. Cep55 promotes cytokinesis of neural progenitors but is dispensable for most mammalian cell divisions. Nat Commun 11. doi:10.1038/s41467-020-15359-w

Thoresen SB, Campsteijn C, Vietri M, Schink KO, Liestøl K, Andersen JS, Raiborg C, Stenmark H. 2014. ANCHR mediates Aurora-B-dependent abscission checkpoint control through retention of VPS4. Nat Cell Biol 16:550–560. doi:10.1038/ncb2959

Tripathi K, Parnaik VK. 2008. Differential dynamics of splicing factor SC35 during the cell cycle. J Biosci 33:345–354. doi:10.1007/s12038-008-0054-3

Turner BM, Franchi L. 1987. Identification of protein antigens associated with the nuclear matrix and with clusters of interchromatin granules in both interphase and mitotic cells. J Cell Sci 87:269–282.

Umbreit NT, Zhang CZ, Lynch LD, Blaine LJ, Cheng AM, Tourdot R, Sun L, Almubarak HF, Judge K, Mitchell TJ, Spektor A, Pellman D. 2020. Mechanisms generating cancer genome complexity from a single cell division error. Science. 368. doi:10.1126/science.aba0712

Uversky VN. 2017. Intrinsically disordered proteins in overcrowded milieu: membrane-less organelles, phase separation, and intrinsic disorder. Curr Opin Struct Biol 44:18–30. doi:10.1016/j.sbi.2016.10.015

Vietri M, Radulovic M, Stenmark H. 2020. The many functions of ESCRTs. Nat Rev Mol Cell Biol 21:25–42. doi:10.1038/s41580-019-0177-4

Yasui Y, Urano T, Kawajiri A, Nagata KI, Tatsuka M, Saya H, Furukawa K, Takahashi T, Izawa I, Inagaki M. 2004. Autophosphorylation of a newly identified site of Aurora-B is indispensable for cytokinesis. JBC 279:12997–13003. doi:10.1074/jbc.M311128200

Zhai Q, Landesman MB, Chung H-Y, Dierkers A, Jeffries CM, Trewhella J, Hill CP, Sundquist WI. 2011. Activation of the retroviral budding factor ALIX. J Virol 85:9222–9226. doi:10.1128/jvi.02653-10

