## Supplementary Information for "Abscission Checkpoint Bodies Reveal a New Facet of Abscission Checkpoint Control"

**This PDF file includes:**

Figure Supplements 1S1 to 6S3
Tables S1 to S2

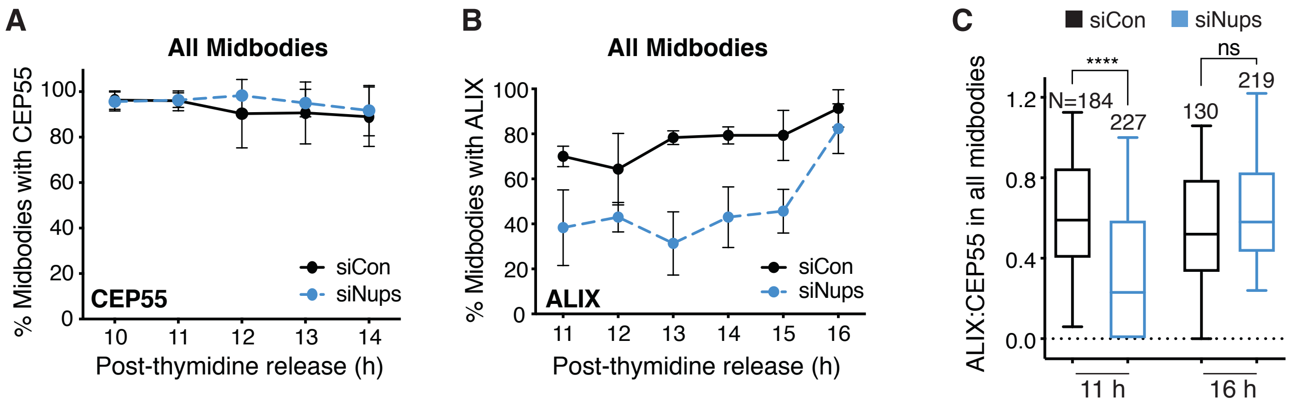


**Figure 2—figure supplement 1.** The abscission checkpoint delays ALIX recruitment in the total midbody population. (**A-B**) Timecourse quantification of CEP55 (**A**) or ALIX **(B**) recruitment to midbodies in control (siControl) and checkpoint-activated (siNup153/50) cells. N=300 midbodies scored (total for each timepoint), n=3 biological replicates. (**C**) Quantification of ALIX:CEP55 relative intensity at individual midbodies in control and checkpoint-activated cells from n=2 biological replicates. Throughout figure supplements: experimental timeline refers to Figure 1B unless otherwise noted. Line and bar graphs represent mean ± standard deviation unless noted. Boxplots represent the 25$^{th}$, median, and 75$^{th}$ percentile of values. Whiskers represent the 10$^{th}$ and 90$^{th}$ percentiles. P<0.05:*, P≤0.01:**, P≤0.001***, P≤0.0001****. See Methods for statistical tests used. N=total from all replicates.

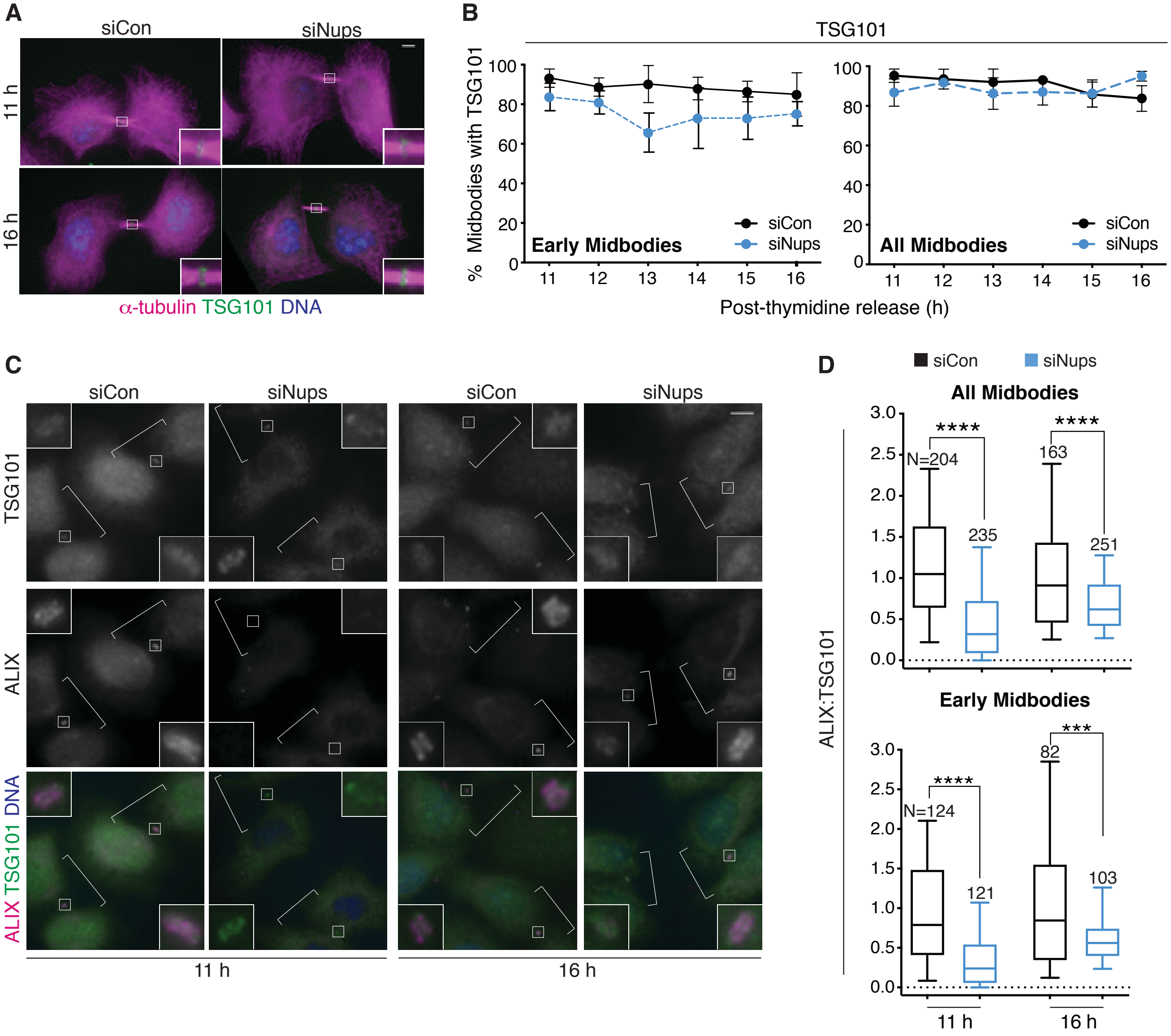


**Figure 2—figure supplement 2**. The abscission checkpoint does not delay TSG101/ESCRT-I recruitment to the midbody. (**A-B**) Immunofluorescence and timecourse quantification of TSG101 recruitment to midbodies in control and checkpoint-activated cells. N=400 midbodies scored/timepoint from n=4 biological replicates. (**C-D**) Immunofluorescence and quantification of ALIX:TSG101 relative intensity at individual midbodies in control and checkpoint-activated cells from n=2 biological replicates. Throughout figure supplements: White brackets mark cells confirmed to be midbody-stage by α-tubulin, staining not shown. Scale bars are 5 μm unless noted. Data points without visible error bars (as in B, "All Midbodies" siCon 14 h) have SDs that do not extend outside the data point.

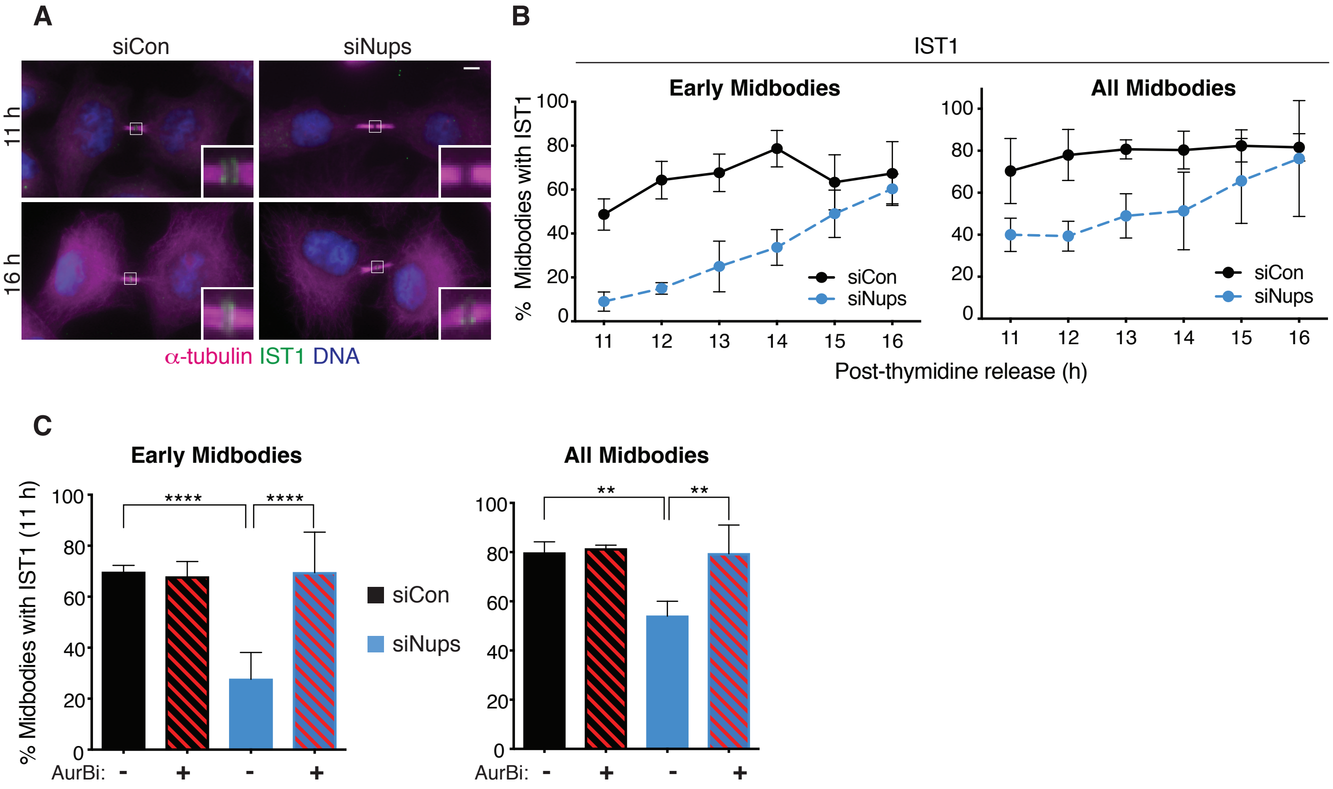


**Figure 2—figure supplement 3.** The abscission checkpoint delays IST1 recruitment to the midbody. (**A-B**) Immunofluorescence and timecourse quantification of IST1 recruitment to midbodies in control and checkpoint-activated midbodies. N=300 midbodies scored/timepoint from n=3 biological replicates. (**C**) Quantification of IST1 recruitment to early and total midbodies, 11 hours post-thymidine release, with/without checkpoint activation, and with/without 30 minutes AurBi added at 10.5 h. N=300 midbodies scored/condition from n=3 biological replicates.

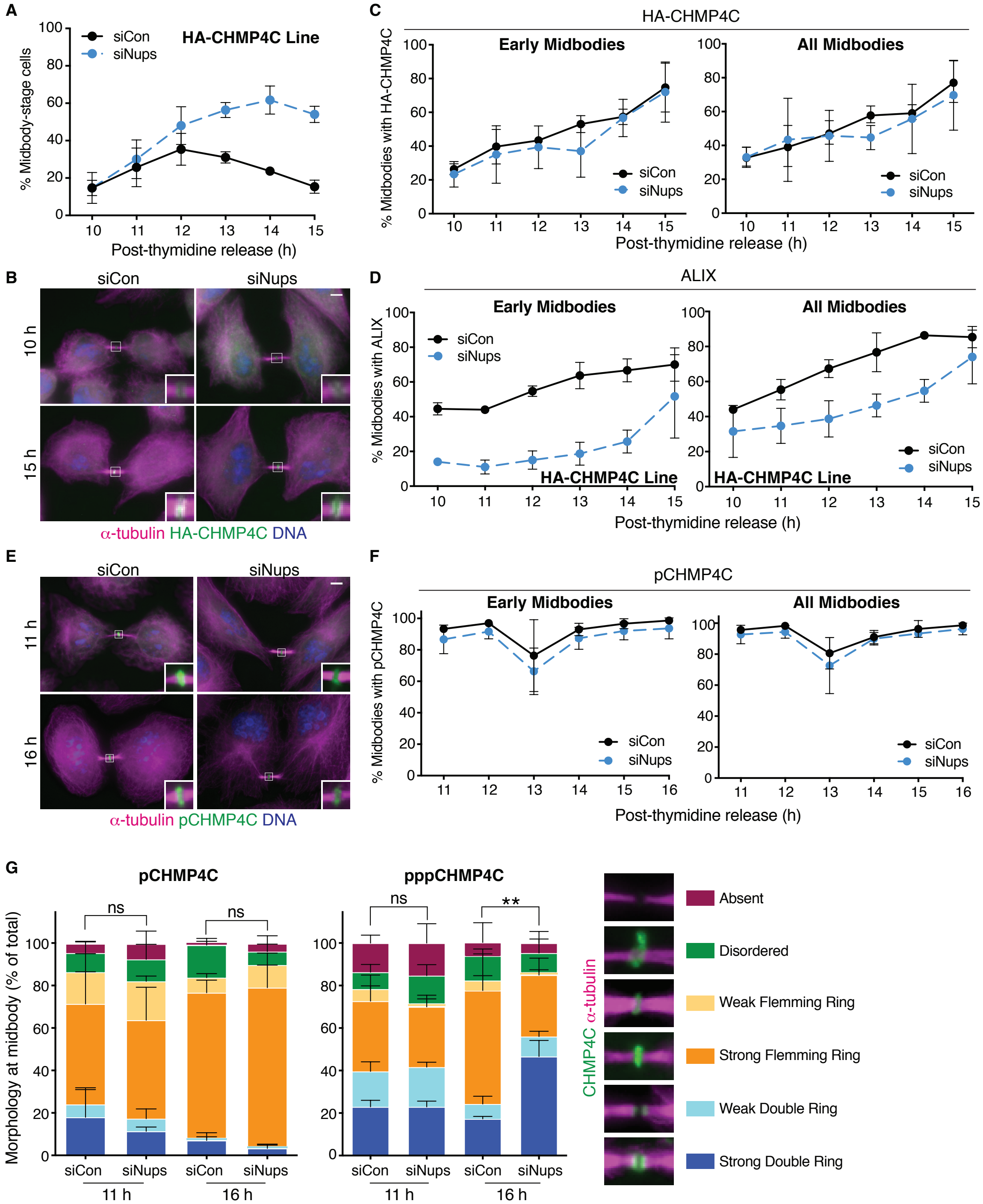


Figure 3—figure supplement 1. The abscission checkpoint does not delay CHMP4C recruitment to the midbody. (A-C) Midbody quantification (A), immunofluorescence (B), and timecourse quantification of HA-CHMP4C recruitment to midbodies (C) in control and checkpoint-activated cells, using a HeLa cell line stably expressing HA-CHMP4C. (A) N=950 cells scored/timepoint from n=3 biological replicates (C) N=300 midbodies scored/timepoint from n=3 biological replicates. (D) Timecourse quantification of ALIX recruitment to midbodies in control and checkpoint-activated HA-CHMP4C-expressing cells. N=300 midbodies scored/timepoint, n=3 biological replicates. (E-F) Immunofluorescence and timecourse quantification of pCHMP4C recruitment to midbodies in control and checkpoint-activated HeLa cells. N=300 midbodies/timepoint, n=3 biological replicates. (G) Quantification of p/pppCHMP4C morphology at the midbody in control and checkpoint-activated cells 11 h and 16 h post-thymidine release. N=300 midbodies/condition, n=3 biological replicates. P-values compare combined weak and strong double rings.

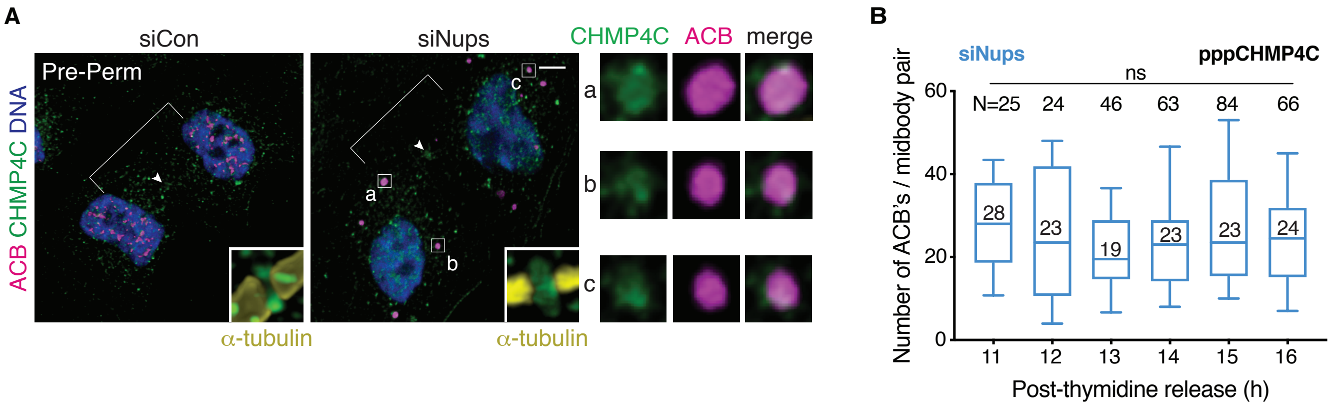


**Figure 3—figure supplement 2.** CHMP4C is detected in ACBs, which are maintained in stable numbers when the abscission checkpoint is active**.** (**A**) Confocal z-projections of pre-permeabilized midbody-stage cells under asynchronous (48 h) control and checkpoint-activated conditions co-stained for CHMP4C and an ACB marker (pSC35, see Figure 4). (**B**) Quantification of number of ACBs, tracked with α-pppCHMP4C, per midbody-stage cell after checkpoint-activation by siNups treatment. n=3 biological replicates; the median ACB number is indicated. Throughout figure supplements: White arrowhead marks Flemming body where detectable. Flemming body insets from confocal imaging are 2.3 μm wide and are enhanced for brightness. ACBs have a brighter signal than the Flemming body, thus some Flemming bodies are seen only in enhanced inset. ACB enlargements are 2 μm wide.

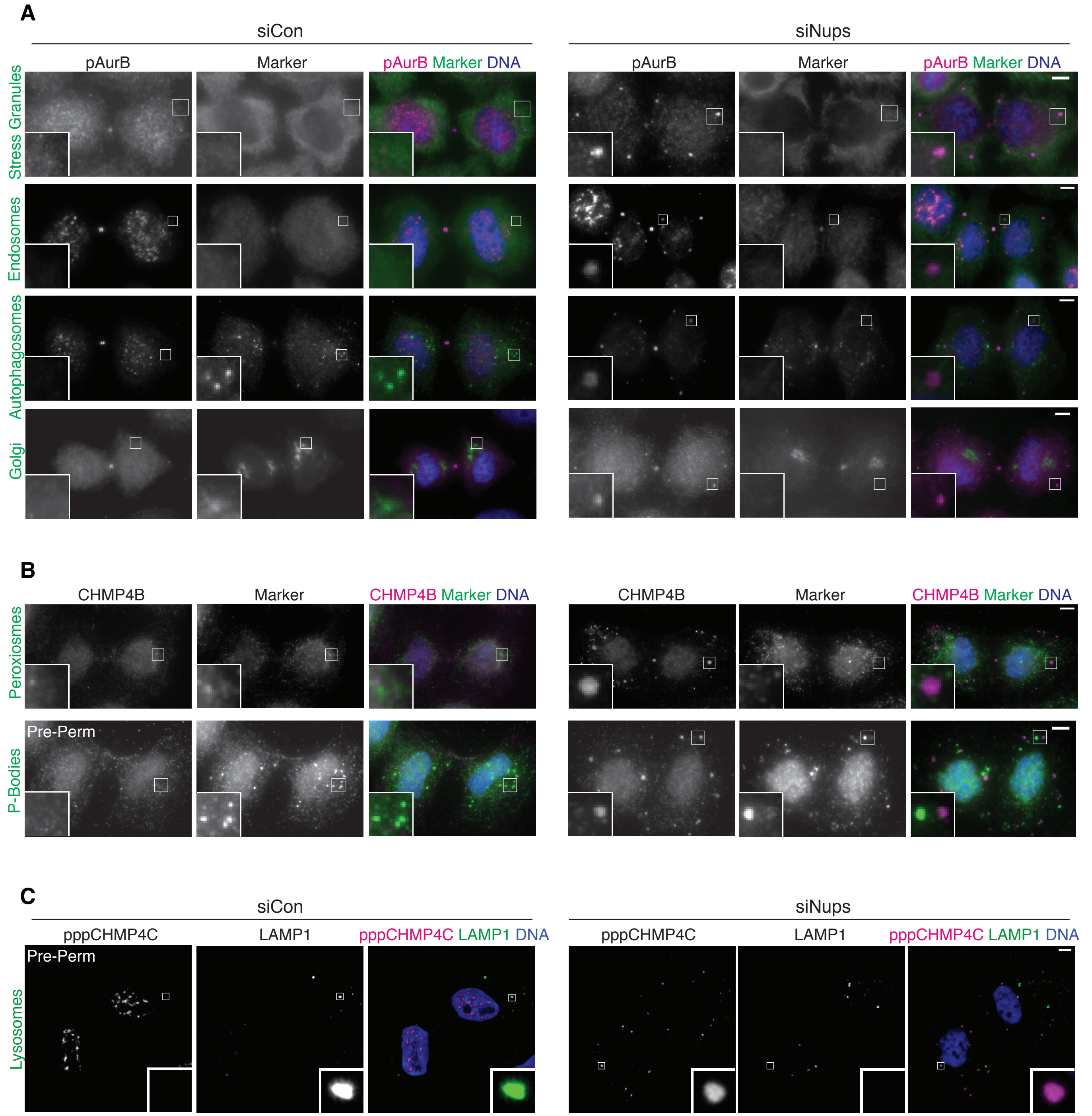


**Figure 4—figure supplement 1.** Abscission checkpoint bodies do not colocalize with a variety of subcellular organelles and structures. (**A-B**) Immunofluorescence of asynchronous (48 h) control and checkpoint-activated (siNups) midbody-stage cells stained to detect ACBs (marked by (**A**) α-pAurB or (**B**) α-CHMP4B), and specific organelle/substructure markers simultaneously. Antibodies used were as follows; stress granules:α-G3BP1, early endosomes:α-EEA1, autophagosomes:α-LC3ß, Golgi:α-giantin, peroxisomes:α-PEX14, P-bodies:α-DDX6. (**C**) Confocal z-projections of pre-permeabilized asynchronous (48 h) control and checkpoint-activated midbody-stage cells co-stained for ACBs (marked by α-pppCHMP4C) and lysosomes (marked by α-LAMP1).

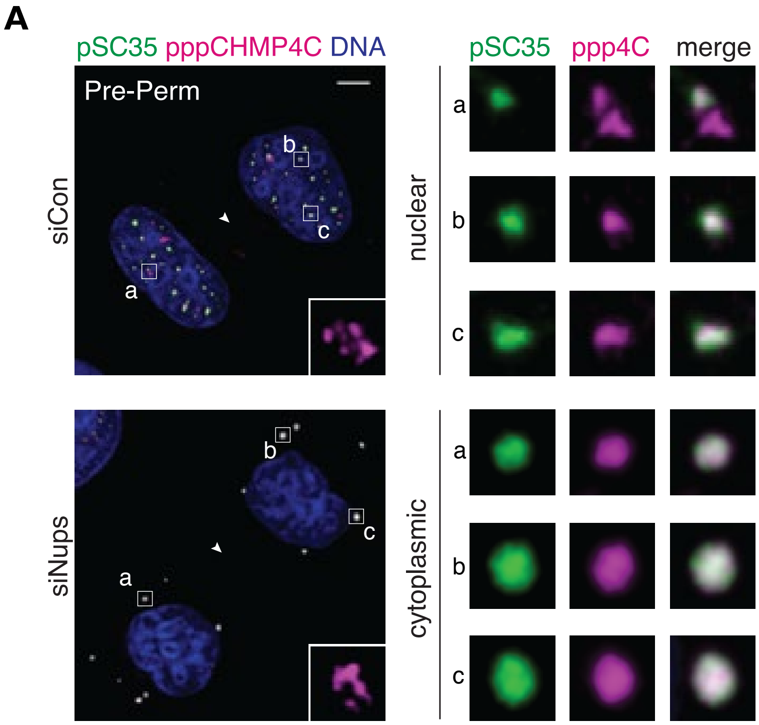


**Figure 4—figure supplement 2.** Colocalization of ACB components. (**A**) Confocal z-projections of pre-permeabilized midbody-stage cells under asynchronous (48 h) control and checkpoint-activated conditions, co-stained for ACB components as indicated.

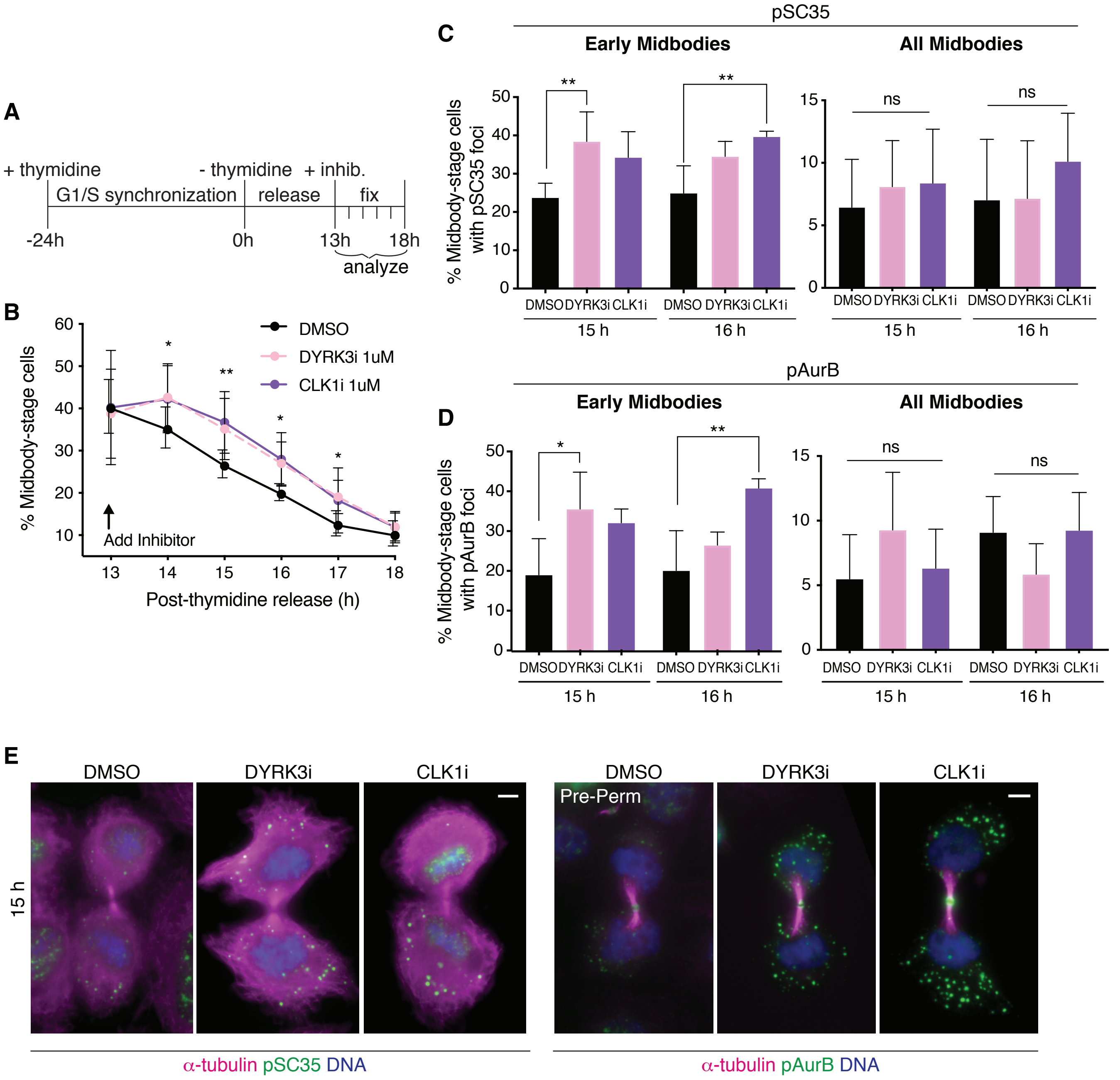


**Figure 4—figure supplement 3.** Interfering with timely resolution of MIGs delays cytokinetic abscission. (**A**) Time schematic of fixed-imaging experiments after treatment with DMSO, 1 µm DYRK3i, or 1 µm CLK1i. Inhibitors were added 13 hours post-thymidine release after most cells had completed metaphase to avoid a confounding metaphase arrest. (**B**) Quantification of midbody-stage cells after treatment as diagrammed in (**A**). N=1200 cells scored/timepoint from n=4 biological replicates. (**C-E**) Quantification and immunofluorescence of midbody-stage cells with cytoplasmic foci marked by α-pSC35 or α-pAurB treated as diagrammed in A. N=400 midbody-stage cells scored/condition from n=4 biological replicates.

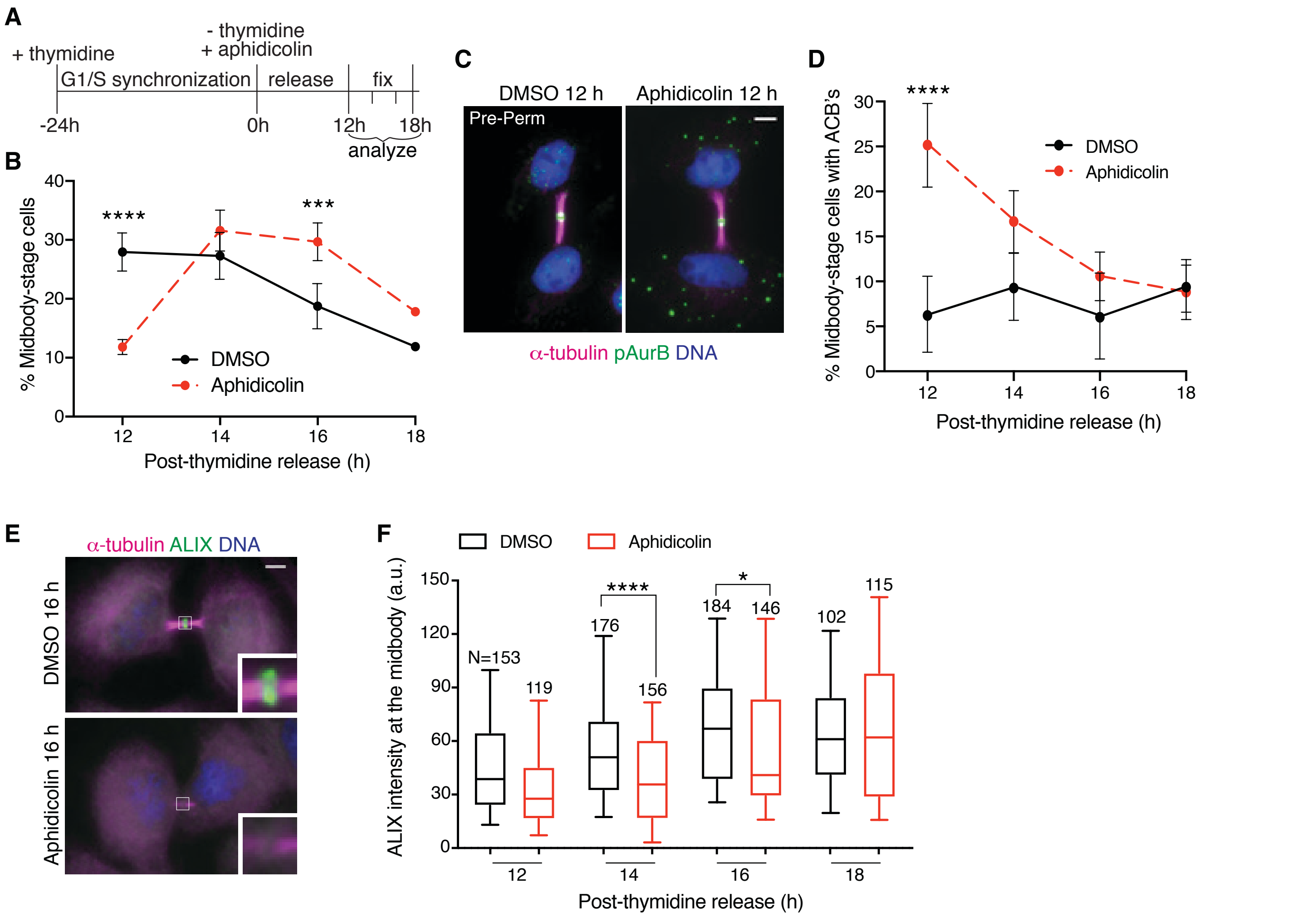


**Figure 5—figure supplement 1.** Abscission checkpoint bodies and ALIX recruitment delay are triggered by replication stress. (**A-B**) Timeline schematic (**A**) and quantification (**B**) of synchronized midbody-stage cells with/without 0.4 μM aphidicolin treatment following thymidine release. N=900 cells scored/timepoint from n=3 biological replicates. (**C-D**) Immunofluorescence and quantification of midbody-stage cells with ACB's after treatment as in (**A**). N=300 midbody-stage cells scored/timepoint from n=3 biological replicates. (**E-F**) Immunofluorescence and quantification of ALIX intensity at midbodies after treatment as in (**A**). n=3 biological replicates.

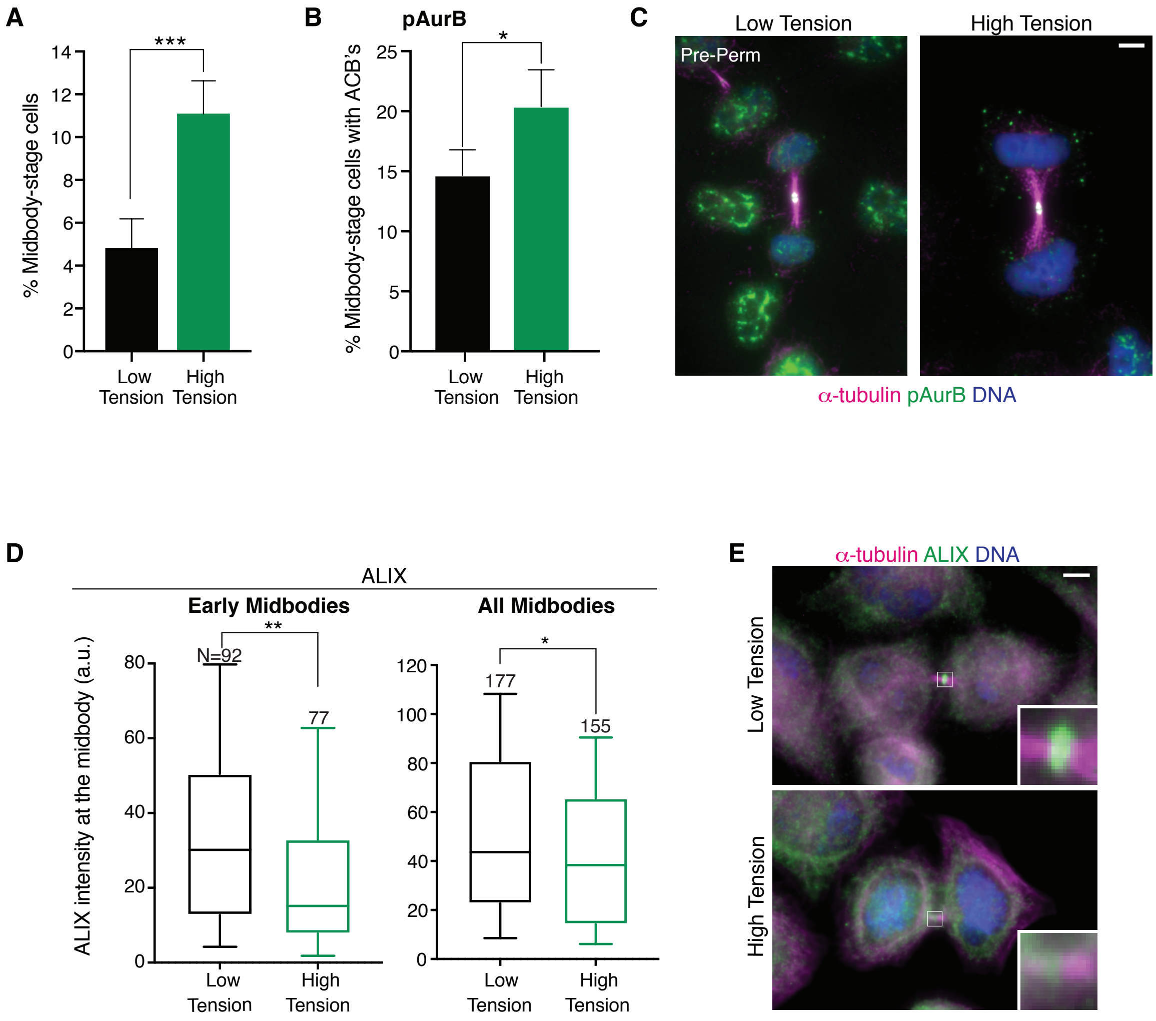


**Figure 5—figure supplement 2.** Abscission checkpoint bodies and ALIX recruitment delay are triggered by intercellular tension. (**A**) Quantification of midbody-stage cells plated at high density (low tension) or low density (high tension). N=1200 cells scored/condition from n=4 biological replicates. (**B-C**) Quantification and immunofluorescence of midbody-stage cells with ACB's marked by α-pAurB when plated at high or low density. N=400 cells scored/condition from n=4 biological replicates. (**D-E**) Quantification and immunofluorescence of ALIX intensity at midbodies in cells plated at high or low density. n=4 biological replicates.

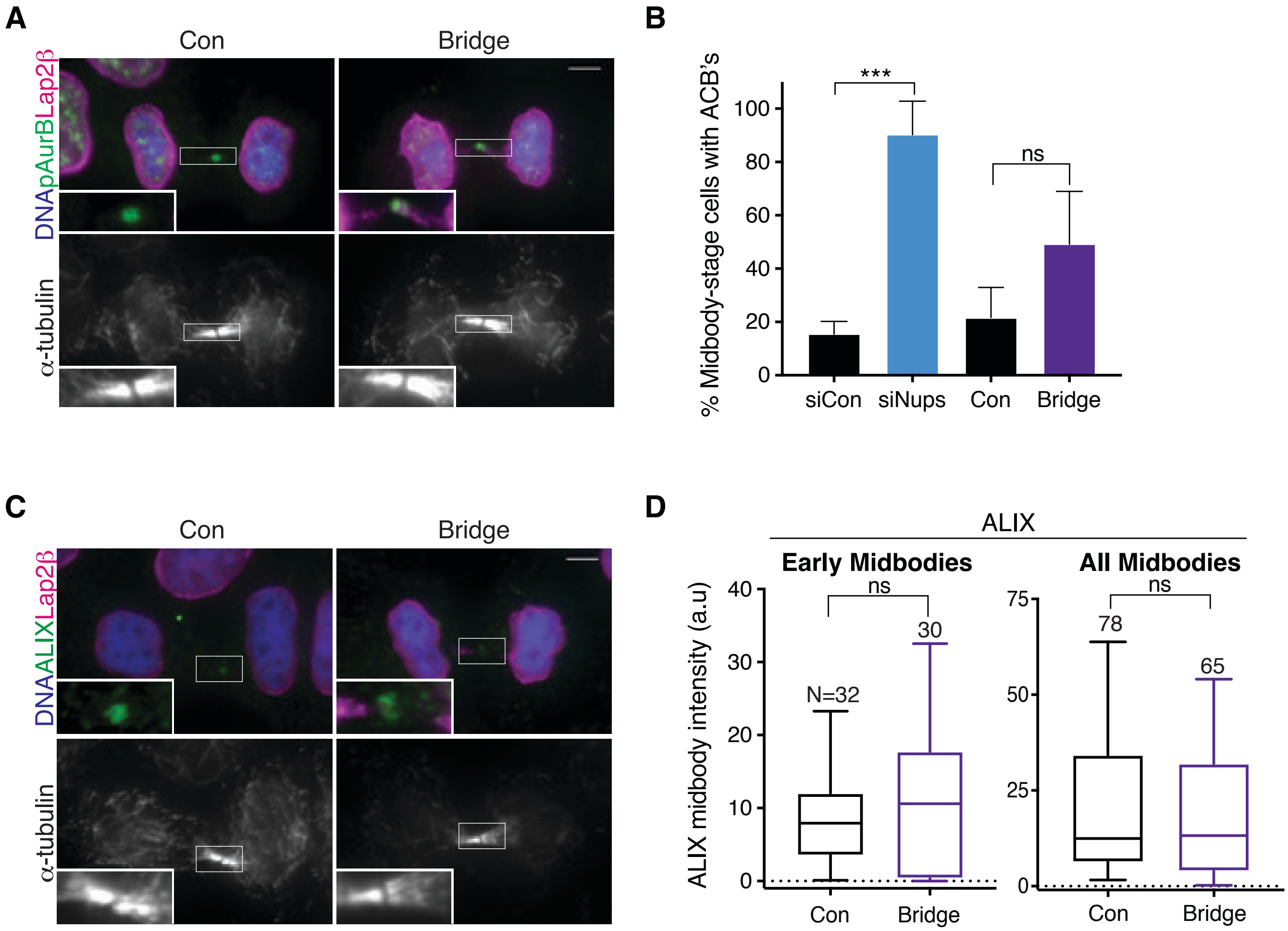


**Figure 5—figure supplement 3.** Chromatin bridges do not significantly trigger abscission checkpoint bodies or ALIX recruitment delay. (**A**) Immunofluorescence of representative cells with and without chromatin bridges. (**B**) Quantification of midbody-stage cells with ACBs both with/without siNups treatment (for comparison) and with/without lagging chromatin bridges. Rare, naturally occurring errors were scored, accounting for the small sample size: N≥142 midbodies scored/condition from n=3 biological replicates. Note that there was a trend toward increased percentages of cells with ACBs when bridges were present, but it did not reach statistical significance. (**C-D**) Representative immunofluorescence and quantification of ALIX recruitment to midbodies in cells with and without chromatin bridges. Insets in (**A and C**) were enhanced for brightness. n=4 biological replicates.

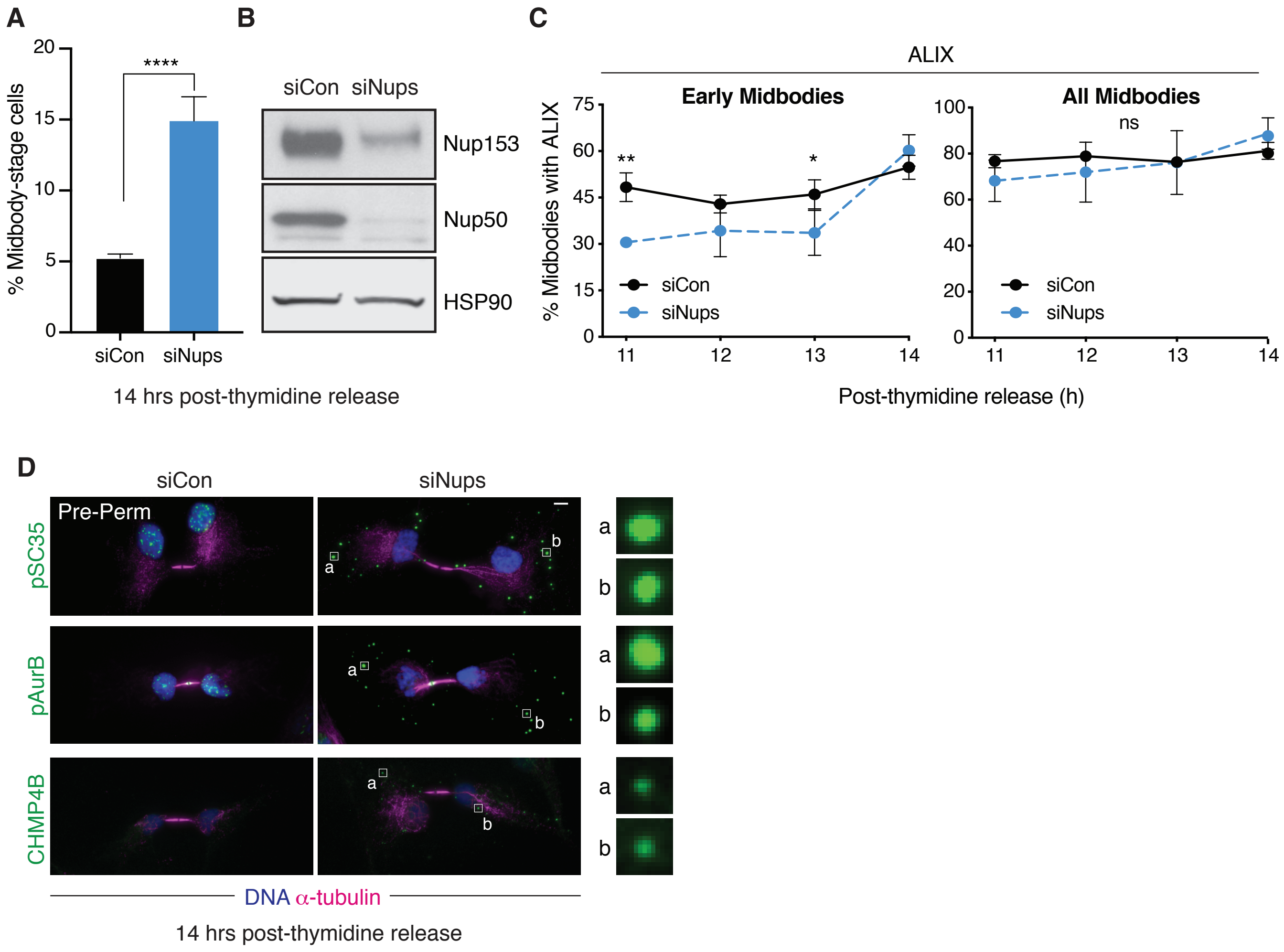


**Figure 5—figure supplement 4.** RPE1 cells display hallmarks of the abscission checkpoint, including ALIX recruitment delay and ACB formation. (**A-B**) Quantification of midbody-stage cells and western blot of lysates from RPE1 cells in control and checkpoint-activated conditions 14 h following thymidine release. N=800 cells scored/condition from n=3 biological replicates. (**C**) Timecourse quantification of ALIX recruitment to midbodies in control and checkpoint-activated cells. N=250 midbodies scored/timepoint from n=3 biological replicates. (**D**) Immunofluorescence of pre-permeabilized control and checkpoint-activated cells 14 h following thymidine release, stained for ACB markers as indicated.

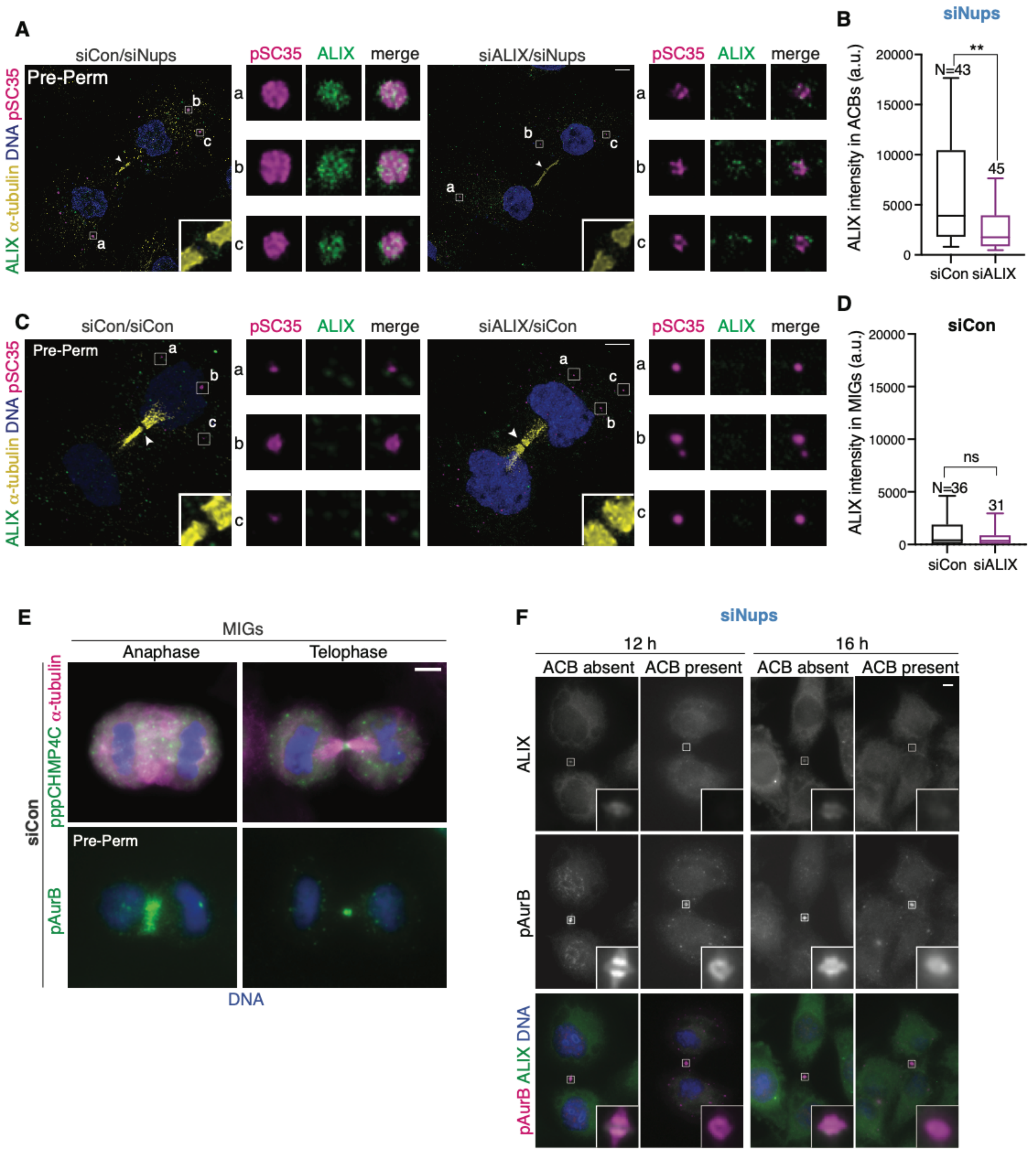


Figure 6—figure supplement 1. ALIX localizes specifically to ACBs. (A-B) Confocal z-projections of pre-permeabilized cells (A) and quantification (B) of ALIX brightness/area in ACBs under asynchronous conditions (72 h after transfection with siCon/siNups or siALIX/siNups). ACB area was defined by α-pSC35. n=3 biological replicates. (C-D) Confocal z-projections of pre-permeabilized cells (C) and quantification (D) of ALIX brightness/area in MIGs under asynchronous conditions (72 h after transfection with siCon/siCon or siALIX/siCon). MIG area was defined by α-pSC35. n=3 biological replicates (E) Representative images of MIGs marked by α-pppCHMP4C or α-pAurB. (F) Representative images from quantification in Fig. 6E using α-pAurB to mark ACBs.

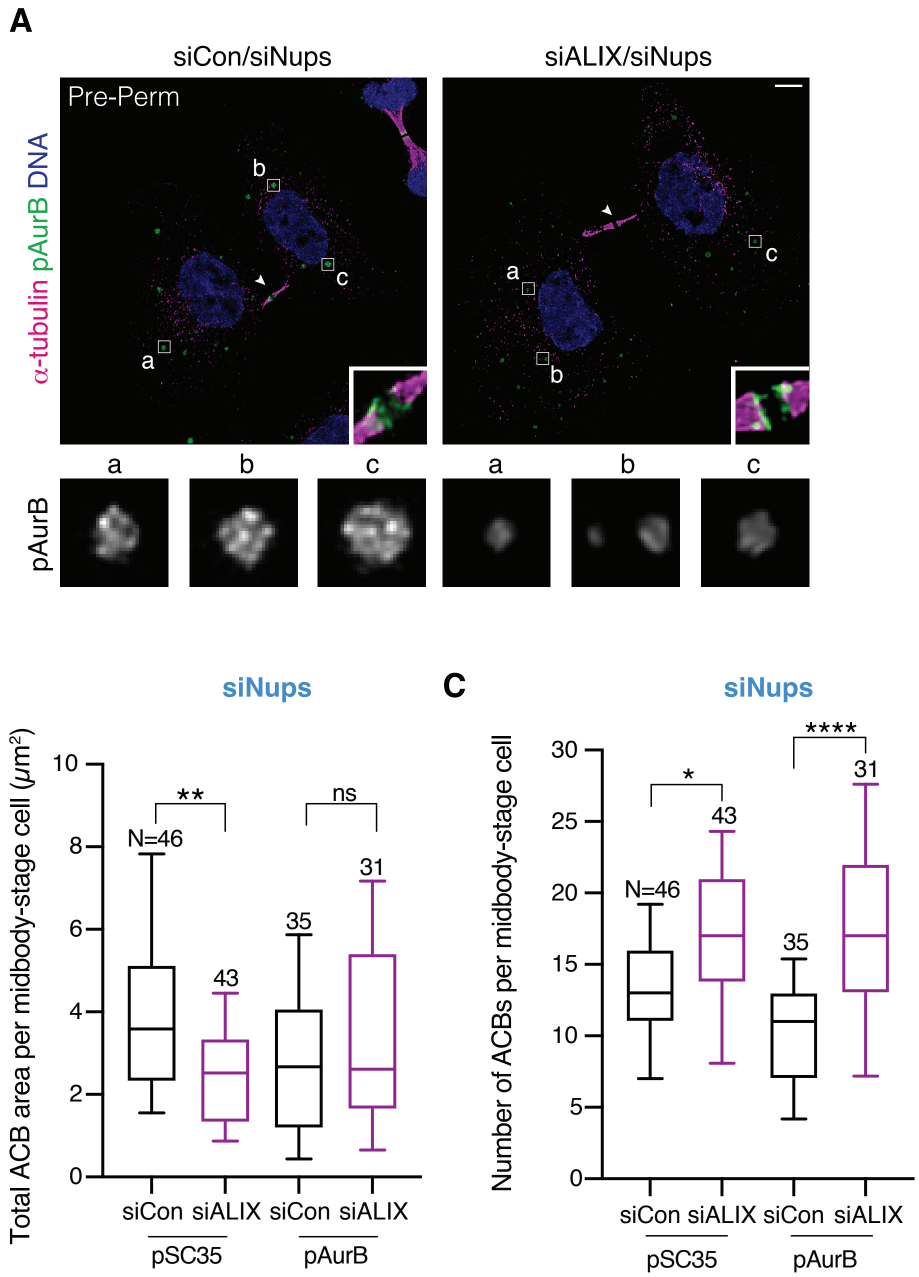


**Figure 6—figure supplement 2.** ACB size is dependent on ALIX. (**A**) Confocal z-projections of pre-permeabilized cells and quantification of ACBs in asynchronous conditions (72 h after treatment with combinations of siCon, siNups, or siALIX as indicated). ACBs were marked by α-pAurB or α-pSC35. (**B-C**) Quantification of total ACB area/midbody-stage cell (**B**) and number of ACBs/midbody-stage cell (**C**) after treatment as in **A.** n=3 biological replicates.

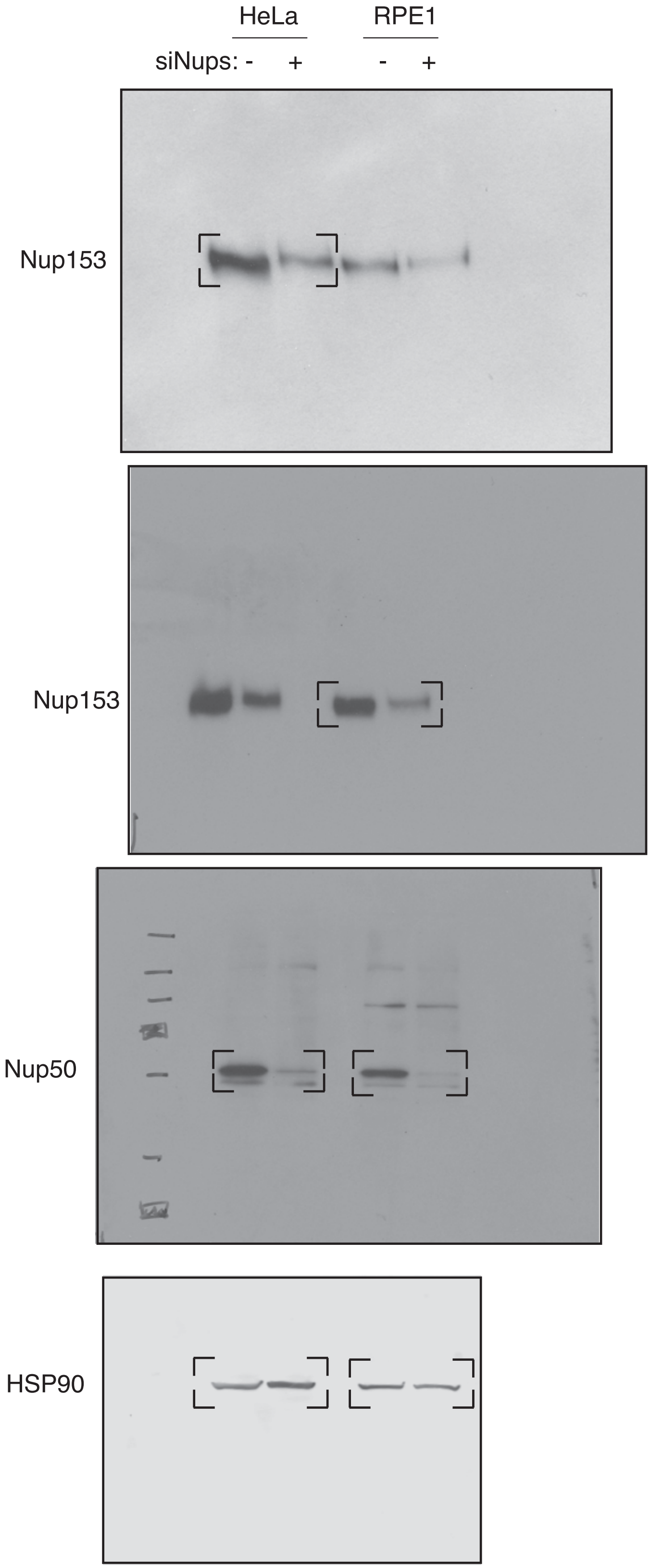


**Figure 6—figure supplement 3.** Uncropped western blots. **(A)** Uncropped western blots of Figure 1B and Figure 5—figure supplement 4B.

**Table S1. Antibodies**
Antibodies used for western blot/immunofluorescence in this study.

| Protein | Host Organism | Dilution | Conditions | Source | Product # (If Applicable) |
|---|---|---|---|---|---|
| $\alpha$-tubulin | Rabbit | IF: 1:1000 | Fix: Methanol, PFA, PHEM | Abcam | ab18251 |
| $\alpha$-tubulin | Rat | IF: 1:1000 | Fix: Methanol, PFA, PHEM | Accurate Chem. & Scientific Corp. | YSRTMCA77P |
| $\alpha$-tubulin | Chicken | IF: 1:400 | Fix: PFA, PHEM | Abcam | Ab89984 |
| $\alpha$-tubulin | Chicken | IF: 1:1500 | Fix: Met, PFA, PHEM | Synaptic Systems | 104 302 |
| ALIX | Rabbit | IF: 1:500<br>WB: 1:1000 | Fix: Methanol, PFA, PHEM<br>WB: PVDF | Covance | UT375 |
| ALIX 2H12 | Mouse | IF: 1:250 | Fix: Methanol, PHEM | Santa Cruz | sc-53539 |
| CEP55 | Rabbit | IF: 1:1000 | Fix: -20 Methanol 10 min, $NH_4Cl$ 15 min, 0.2% Triton-X 7 min | Francis Barr<br>(Bastos and Barr, 2010) | N/A |
| CHMP4B | Rat | IF: 1:500 | Fix: PFA, PHEM | Covance | UT757 |
| CHMP4C | Rabbit | IF: 1:100 | Fix: PHEM | Abcam | ab155668, |
| DDX6 | Rabbit | IF: 1:400 | Fix: PHEM | Bethyl Laboratories | A300-461A |
| EEA1 | Mouse | IF: 1:500 | Fix: Methanol | Santa Cruz | sc-137130 |
| G3BP1 | Mouse | IF: 1:500 | Fix: Methanol | Santa Cruz | sc-365338 |
| Giantin | Mouse | IF: 1:1000 | Fix: Methanol | Abcam | ab37266 |
| HA | Mouse | IF: 1:400 | Fix: Methanol | Cell Signaling | 2367 |
| HSP90 | Mouse | WB: 1:10,000 | WB: PVDF | Santa Cruz | sc-13119 |
| IST1 | Rabbit | IF: 1:500 | Fix: Methanol | Covance<br>(Affinity-Purified) | UT560 |
| LAMP1 | Mouse | IF: 1:500 | Fix: Methanol, PHEM | Santa Cruz | sc-18821 |
| Lap2ß | Mouse | IF: 1:100 | Fix: Methanol, PHEM | Brian Burke | N/A |
| LC3ß | Mouse | IF: 1:250 | Fix: Methanol | Santa Cruz | sc-271625 |
| Nup153 | Mouse | WB: 1:50 | WB: PVDF, 6% Gel | Brian Burke | SA1 |
| Nup50 | Rabbit | WB: 1:500 | WB: PVDF<br>Not suitable for IF | Katharine Ullman<br>(Mackay et al., 2010) | N/A |
| pAurB | Rabbit | IF: 1:500 | Fix: Methanol, PHEM | Rockland | 600-401-677 |

| | | | | | |
|---|---|---|---|---|---|
| pCHMP4C | Rabbit | IF: 1:500 | Fix: Methanol | Pier Paulo D'avino (Capalbo et al., 2016)) | N/A |
| pppCHMP4C | Rabbit | IF: 1:500 | Fix: Methanol, PHEM | Pier Paulo D'avino (Capalbo et al., 2016) | N/A |
| PEX14 | Rabbit | IF: 1:500 | Fix: PFA | Millipore | ABC142 |
| pSC35 | Mouse | IF: 1:2000 | Fix: Methanol, PHEM | Abcam | ab11826 |
| SR domain | Mouse | IF: 1:250 | Fix: Methanol, PHEM | Sant Cruz | sc-13509 |
| TSG101 | Mouse | IF: 1:500 | Fix: -20 Methanol 20 min, 0.05% Triton-X 10 min | Genetex | GTX70255 |

**Table S2. siRNAs**
siRNAs used in this study.

| Protein Target | Sense Sequence | Concentration | Min. Length of Knockdown | Reference |
|---|---|---|---|---|
| ALIX | CCUGGAUAAUGAUGAAGGA | 20-30 nm | 72 hours | (Morita et al., 2007) |
| CHMP4C | CACUCAGAUUGAUGGCACA | 100 nm | 72 hours | (Sadler et al., 2018) |
| Control | GCAAAUCUCCGAUCGUAGA | | 42 hours | (Mackay et al., 2010) |
| Nup153 | GGACUUGUUAGAUCUAGUU | 10 nm | 42 hours | (Mackay et al., 2010) |
| Nup50 | GGAGGACGCUUUUCUGGAU | 10 nm | 42 hours | (Mackay et al., 2010) |
